# Reversible Opto-Chemical activation of KRASG12V signaling with near single-cell precision

**DOI:** 10.64898/2026.06.29.735036

**Authors:** Luis Manuel Muñoz-Nava, Maria Bespalova, Mario O. Caracci, Kitso Ata Kewagamang, Birga Soetje, Sabrina Seidler, Kirsten Michel, Holger Vogel, Janicke Maerz, Philippe I. H. Bastiaens

**Affiliations:** Department of Systemic Cell Biology, Max Planck Institute of Molecular Physiology, Dortmund, Germany; Faculty of Chemistry and Chemical Biology, TU Dortmund, Dortmund, Germany

**Keywords:** oncogenic KRAS, acute signaling, optogenetics, chemically inducible dimerization, mouse intestinal organoids

## Abstract

*KRAS* mutations drive some of the most lethal carcinomas, and genomic and inducible systems have established many of the cellular and tissue-level consequences. However, these approaches operate at the level of oncogene expression, allowing for cellular adaptation that masks the individual role of KRAS oncoprotein signaling. Here, we developed a reversible Opto-Chemical system to activate KRAS signaling by chemically translocating a cytosolic mutant KRASG12V G-domain to the plasma membrane upon light or small-molecule input. In MDCK cells, the G-domain plasma membrane recruitment activated downstream signaling and reduced collective migration. In mouse small Intestinal Organoids, G-domain recruitment promoted increased crypt size and number under Epidermal Growth Factor-deprived conditions. We further showed that the increased number of crypts depended on continuous KRASG12V signaling. Finally, under the same deprived conditions, localized activation in just one budding crypt promoted crypt formation compared to controls. This system decouples oncoprotein activity from oncogene expression, allowing to investigate the KRAS signaling contribution to early epithelial transformation.

## Introduction

*KRAS* point mutations are well-known drivers for some of the most lethal human carcinomas, namely pancreatic ductal adenocarcinoma, colorectal carcinoma, and lung adenocarcinoma (Cox *et al*, 2014; Moore *et al*, 2020). KRAS is a small GTPase that actively localizes at the plasma membrane (PM) via a C-terminal anchor (Chandra *et al*, 2012; Zimmermann *et al*, 2013; Schmick *et al*, 2014, 2015) and alternates between active guanosine triphosphate (GTP)-bound and inactive guanosine diphosphate (GDP)-bound states, regulated by guanine nucleotide exchange factors (GEFs) and GTPase-activating proteins (GAPs), respectively (Aronheim *et al*, 1994; Bos *et al*, 2007; Bandaru *et al*, 2019). Once GTP-bound (active), KRAS recruits effector proteins, such as RAF, PI3K, and RalGDS, to the PM (Terai & Matsuda, 2005; Rodriguez-Viciana *et al*, 1994; Mozzarelli *et al*, 2024), reducing their degrees of freedom from three to two dimensions and thereby enhancing their local interactions with proteins and lipids as part of the signal transduction mechanism (Zamir *et al*, 2013; Schmick & Bastiaens, 2014).

*KRAS* mutations, typically at codons 12, 13, or 61, disable GTP-GDP cycling and lock KRAS in a constitutively active GTP-bound state, leading to persistent effector recruitment and uncontrolled downstream signaling (Cox *et al*, 2014; Simanshu *et al*, 2017). Genomic and inducible expression systems have defined many of the cellular and tissue-level consequences of mutant *KRAS* in tumorigenesis (Jinesh *et al*, 2018; Moore *et al*, 2020). These studies have shown that the outcome of *KRAS* mutations is highly context-dependent, and among others, shaped by the presence or absence of additional genetic or epigenetic defects, such as loss of tumor suppressors *APC*, *p53,* or *LKB1* (Sansom *et al*, 2006; Cox *et al*, 2014; Iovanna & Dusetti, 2026). In vitro systems such as mouse Small Intestinal Organoids (mSIOs, Sato *et al*, 2009) offer an interesting model for studying intestinal carcinogenesis associated with *KRAS* activating mutations. For instance, endogenous KRASG12D expression was shown to increase cell proliferation, leading to bigger organoids and increased crypt number compared to controls (Kotani *et al*, 2021). In contrast, inducible overexpression of KRASG12V in mSIOs was reported not to modify organoid development, but led to a modest increase in MAPK activity (Brandt *et al*, 2019). However, these approaches couple oncogene expression to oncoprotein signaling activity within all the cells of the tissue, allowing for cellular adaptations to oncogene expression as shown for pancreatic cells (Mathison *et al*, 2021), making it difficult to investigate the local and individual role of KRAS signaling in tissue transformation (Jonkers & Berns, 2002; Voß *et al*, 2015). Furthermore, a reversible strategy to acutely start as well as to stop KRAS oncoprotein signaling would allow to determine whether early transformation is dependent on continuous oncoprotein signaling (Weinstein, 2002; Pagliarini *et al*, 2015) or can be undone.

Cellular optogenetics and chemical-inducible systems (CID) can, in principle, address this challenge by enabling acute and spatially controlled activation of signaling nodes. A split-small GTPase system based on CID has been reported to successfully control KRAS, Cdc42, and RhoA signaling in mammalian cells by translocation of the GTPase catalytic domain to the PM (He *et al*, 2024, 2025). In addition, optogenetic tools targeting proteins both upstream and downstream of KRAS, including SOS RAS-GEF, RAF, PI3K, Akt, and MEK, have been developed (Toettcher *et al*, 2011, 2013; Wend *et al*, 2014; Katsura *et al*, 2015; Krishnamurthy *et al*, 2016; Johnson *et al*, 2017; Zhou *et al*, 2017; Kramer *et al*, 2021; Gagliardi *et al*, 2023). These approaches have shed some light on signaling dynamics and cell-fate decisions, but they do not fully capture the regulatory context of KRAS oncoprotein signaling activities or lack a reversibility strategy. Activation of isolated downstream proteins bypasses KRAS activity and negative feedback regulation (Lake *et al*, 2016), while translocation of SOS to the PM activates all RAS isoforms in its vicinity, rather than a specific variant (Toettcher *et al*, 2013).

In this paper, we demonstrate a novel approach for acute and reversible activation of KRAS oncoprotein downstream signaling in both 2D and 3D environments, i.e., MDCK cell monolayers and mSIOs, with near single-cell precision, that decouples oncoprotein signaling activity from oncogene expression. The principle combines CID and Opto-Chemical manipulation published earlier (Spencer *et al*, 1993; Fegan *et al*, 2010; Liu *et al*, 2014; Chen & Wu, 2018; Chen *et al*, 2018a), which allows for the reversible KRASG12V G-domain concentration to the PM (Opto-CID-KRASG12V). This mechanism works under the premise that PM-enrichment and dimensionality reduction are essential for KRAS signaling transduction; the expression of cytoplasmic KRASG12V G-domain should not result in oncogenic transformation. Our results showed differences in MDCK cell collective migration and mSIOs morphology during development by concentrating KRASG12V G-domain to the PM.

## Results

### Light-induced plasma membrane localization of KRASG12V triggers reversible oncoprotein signaling

To dissect the oncoprotein signaling activity from oncogene expression with space and time control, we generated the genetically encoded two-part Opto-CID-KRASG12V system (Fig. 1 A), consisting of: 1) CD-GDom, containing the constitutively active G-domain of KRAS bearing the G12V mutation, without the hypervariable region (KRASΔHVR, aa 1-164), fused to a tandem of two *Escherichia coli* dihydrofolate reductase (2XecDHFR; Liu *et al*, 2014; Chen *et al*, 2018a; Chen & Wu, 2018), one mCitrine fluorescent protein and a nuclear exporting signal (NES) to restrict its localization to the cytoplasm (Fig. 1 A, left); dimerized on demand to: 2) TH-tKRAS, which is tethered to the PM via the hypervariable domain of KRAS (HVR) carrying the membrane targeting prenylatable CAAX motif (tKRAS; Liu *et al*, 2014; Chen *et al*, 2018a), fused to a tandem of two modified haloalkane dehalogenase proteins (2XHaloTag; Los *et al*, 2008; Chen *et al*, 2018a) and one mTFP fluorescent protein. The DNA cassette containing both fragments separated by a ribosomal skipping sequence (P2AT2A) was cloned into a single piggyBac vector (Wang *et al*, 2008), enabling the generation of stable cell lines and organoids (Fig. S1 A). Dimerization is achieved by using the photocaged NvocTMP-Cl dimerizer (Fig. 1 A, middle and right), custom synthesized (ChiroBlock GmbH) as previously described (Ballister *et al*, 2014; Chen *et al*, 2018b; Los *et al*, 2008). This dimerizer consists of: 1) NvocTMP, the ecDHFR ligand trimethoprim (TMP), caged with the 6-nitroveratryl carbamate (Nvoc) that can be photo-uncaged by 405 nm light irradiation, resulting in the exposed TMP molecule moiety, and 2) a HaloTag covalent ligand (Cl), chloroalkane group; both moieties were linked through a polyethylene glycol (PEG) linker. When both moieties are exposed, the complex ecDHFR-TMP-Cl-HaloTag can be formed, concentrating the CD-GDom fragment to the PM (Fig. 1 A, middle). Reversibility of CD-GDom localization to the cytoplasm is achieved by adding free TMP, which competes with the TMP moiety of the dimerizer for binding to ecDHFR (Fig. 1A, left and middle).

**Figure 1.**
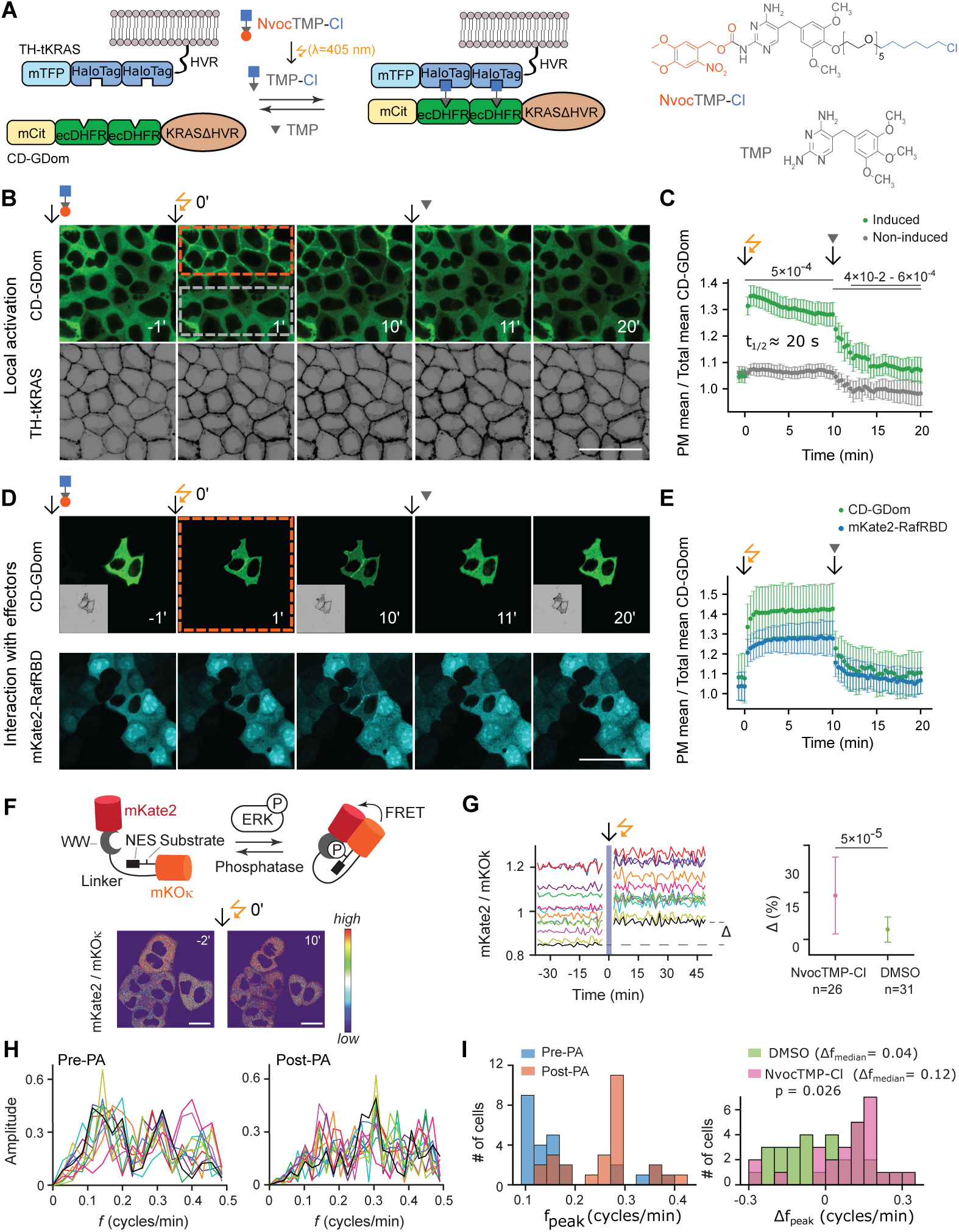
Light-induced space-time control of KRASG12V oncoprotein signaling activity. **(A)** Scheme of the Opto-CID-KRASG12V system, from left to right, PM tethered TH-tKRAS containing mTFP, 2XHaloTag and tKRAS; Cytoplasmic CD-GDom containing mCitrine, 2XecDHFR and KRASΔHVR; NvocTMP-Cl: 6-nitroveratryloxycarbonyl (Nvoc; dark orange ellipse) caging TMP (grey triangle), conjugated to chlorohexyl moiety (Cl; blue square), opto-chemical photo-unccaging (PA) is achieved by 405-nm light irradiation (orange zigzag arrow), TMP molecule for reversibility (left arrow); Stabilization of CD-GDom at the PM after complex ecDHFR-TMP-Cl-HaloTag formation; Chemical Structures of NvocTMP-Cl and TMP. **(B)** Representative micrographs of CD-GDom (green) and TH-tKRAS (inverted grey) fluorescence at the indicated time points in Opto-CID-KRASG12V-MDCK cells after NvocTMP-Cl (10 μM) 1 h incubation, PA via 405-nm light (within the orange dashed rectangle) at time 0 min and the subsequent TMP (10 μM) addition for reversibility after 10 min. **(C)** Scatter plot depicting the ratio of the CD-GDom mean intensity of PM over total (PM + cytoplasm) over time, for the induced (green) and Non-Induced (grey; light grey dashed rectangle in **B**) areas, showing the t_1/2_ time until reaching the maximum PM concentration. *p*-value was calculated by the Mann-Whitney U test with Bonferroni correction. **(D)** Representative micrographs of CD-GDom, TH-tKRAS (inset), and mKate2-RafRBD (cyan) following the same induction and reversibility procedure as in **B** within the complete field of view (orange dotted square). **(E)** Scatter plot depicting the ratio of the CD-GDom mean intensity of PM over total mean intensity for the CD-GDom (green) and mKate2-RafRBD (blue). n = 13 and 12 for **C** and **E**, respectively. Scale bar = 50 μm. **(F, top)** Schematics of Booster-ERK FRET sensor and its operating principle. From the N-terminus, the sensor consists of the acceptor FP mKate2, WW domain, a 244-amino-acid linker, mKOk, ERK substrate peptide, and nuclear export signal (NES), and the donor mKOk. In the phosphorylated state, ERK phosphorylation of the substrate peptide promotes intramolecular binding of the phosphothreonine of the ERK substrate to the WW domain, inducing a conformational change that increases FRET between mKOk and mKate2. **(F, bottom)** Representative mKate2/mKOk ratio micrographs of MDCK cells expressing Opto-CID-KRASG12V-Booster and incubated with NvocTMP-Cl, acquired 2 min before and 10 min after the 405-nm PA. Scale bar, 20 μm. **(G, left)** Corresponding normalized single-cell time courses of the mKate2/mKOk ratio. Each trace was corrected for a global trend and rescaled to its first time point. The blue shaded area indicates PA; Δ denotes the PA-induced change in mKate2/mKOκ ratio, and Δ_mean_ indicates the mean change across all traces shown. **(G, right)** Comparison of PA-induced changes in mKate2/mKOk, Δ, for NvocTMP-Cl-(pink) and DMSO-treated (green) cells. **(H)** FFT amplitude spectra of the normalized mKate2/mKOk ratio oscillations as a function of frequency, *f*, calculated from the pre-PA and post-PA segments of the time courses shown in **G**. Colors were preserved between the time-course and FFT plots for each cell. **(I)** Left: Distributions of peak frequencies, *f*_peak_, derived from pre-PA (blue) and post-PA (orange) FFT spectra for NvocTMP-Cl-treated cells (n=26, from 3 independent cell clusters including cells shown in **H**). Each *f*_peak_ represents the frequency of maximal FFT amplitude from an individual cell’s spectrum. Right: Distributions of per-cell frequency shifts upon PA, defined as Δ*f*_peak_ = *f*_peak_(post-PA) −*f*_peak_(pre-PA), for NvocTMP-Cl (pink) and DMSO-treated (green, control) cells. Δ*f*_median_ indicates median Δ*f*_peak_ values. *p*-value was calculated by the Wilcoxon rank-sum test for **G** and **I**.

We generated an MDCK cell line stably expressing the Opto-CID-KRASG12V system (Fig. S1 B), forming a 2D monolayer that allows for a clear and independent visualization of CD-GDom and TH-tKRAS in the cytoplasm and PM, respectively (Fig. 1 B, left). After incubation with the NvocTMP-Cl dimerizer (10 µM), we photoactivated (PA) a selected group of cells by short-term exposure to 405 nm light in the region of interest (Fig. 1 B, orange dashed line; see Methods), resulting in the translocation of CD-GDom to the PM without a detectable change in non-irradiated cells within the field of view (grey dashed rectangle), demonstrating precise spatio-temporal control over concentration at the PM. The fraction of CD-GDom localized at the PM was quantified as the ratio of the mean fluorescence intensity in the PM masked pixels over that across all pixels (PM + nucleus-cytoplasm; see Methods). This fraction showed a very rapid increase (t_1/2_ ≈ 20 s) after the light irradiation pulse (Fig. 1 C), comparable to the kinetics reported for another opto-chemical system using the NvocTMP-Cl dimerizer (Chen *et al*, 2018a). Subsequent addition of the free TMP molecule in excess (10 µM) at 10 min post-PA rapidly reversed CD-GDom to the cytoplasm, decreasing the PM/total ratio close to baseline levels within 5 min (Fig. 1 B, C).

After we established the acute, reversible, and spatially controlled concentration of CD-GDom to the PM, we asked whether this mechanism was sufficient to induce signaling effectors downstream of KRAS. To this end, we transiently transfected the Opto-CID-KRASG12V construct into MDCK cells stably expressing the RAS Binding Domain (RBD) of c-Raf fused to mKate2 (RafRBD-mKate2). In cells expressing both DNA constructs (Fig. 1 D, CD-GDom+ cells), RafRBD-mKate2 was confined to the cytoplasm, in contrast to non-expressing neighboring cells showing a cytoplasmic-nuclear distribution. This observation is consistent with the GTP-bound state of CD-GDom and the cytoplasmic CD-GDom-RafRBD complex formation. Following incubation with NvocTMP-Cl and a 405 nm light PA pulse (orange dashed rectangle, whole field of view), both proteins rapidly and simultaneously translocated to the PM (Fig. 1 D, E) without affecting the distribution of RafRBD-mKate2 in neighboring cells. Subsequent addition of TMP (10 µM) at 10 min post-PA reverted localization of both CD-GDom and RafRBD-mKate2 to the cytoplasm. These results experimentally illustrate the dimensionality reduction mechanism by which KRAS concentrates effector proteins to the 2D PM (Zamir *et al*, 2013; Schmick & Bastiaens, 2014), here shown by concentrating the CD-GDom-effector complex to the PM in a controlled and reversible manner.

To further confirm the downstream signaling effects of light-induced CD-GDom translocation to the PM, we monitored ERK activity by Förster Resonance Energy Transfer (FRET) using the cytoplasmic Red-Shifted FRET biosensor harbouring the donor mKOk and acceptor mKate2 fluorophores (Booster-ERK, Fig. 1 F; see Methods; Watabe *et al*, 2020). For FRET ratio imaging experiments, we transfected MDCK cells with the Opto-CID-KRASG12V-Booster construct that encoded the Booster-ERK sensor together with the Opto-CID-KRASG12V recruitment module described above (Fig. S1 C-E; see Methods). In both NvocTMP-Cl- (Fig. 1 F-I) and DMSO-incubated cells (Fig. S1 F-H), photoactivation induced a step-like change in the mKate2/mKOk ratio, although the magnitude of the response, Δ, varied markedly between individual cells (Fig. 1 G and Fig. S1 F, down). Moreover, NvocTMP-Cl-treated cells exhibited a larger average Δ than those incubated in DMSO, 14 ± 12 % vs. 3 ± 4 % (Fig. 1 G, right). We next analyzed the structure of mKate2/mKOk oscillations before and after photoactivation for both NvocTMP-Cl- and DMSO-treated cells. In order to do this, each cell’s mKate2/mKOk time course was split into pre-photoactivation (pre-PA) and post-photoactivation (post-PA) segments, and a single-sided FFT amplitude spectrum was calculated for each segment (Fig. 1 H and Fig. S1 G). We then extracted the peak frequency, *f*_peak_, from each spectrum as a frequency corresponding to the maximal spectral amplitude. Comparing *f*_peak_ values between the pre-PA and post-PA segments revealed a clear shift toward higher frequencies in NvocTMP-Cl-treated cells (Fig. 1 I, left), whereas DMSO-treated cells showed no obvious shift (Fig. S1 H). To quantify this change, we calculated the photoactivation-induced shift in peak frequency, Δ*f*_peak_, for each cell as the difference between the post-PA and pre-PA *f*_peak_ values. The distributions of Δ*f*_peak_ for both NvocTMP-Cl and DMSO are shown in Fig. 1 I (right) and reveal a clear shift toward positive frequency changes in NvocTMP-Cl-treated cells, with most values falling between 0 and 0.3 and a median of 0.12. In contrast, DMSO-treated cells displayed a more symmetric distribution around zero, with a median of 0.04. Together, these data indicate that KRASG12V recruitment to the plasma membrane not only increased the Booster-ERK readout, consistent with higher ERK activity, but also altered ERK dynamics towards faster oscillations.

These findings were further supported by an orthogonal, non-photoactivable CID system, based on the SLF-TMP dimerizer promoting ecDHFR and FKBP36V dimerization (Fig. S2 A-D; Clackson *et al*, 1998; Liu *et al*, 2014), in which the KRASG12V G-domain concentrated to the PM after SLF-TMP (2 μM) addition (Fig. S2 E; t_1/2_ = 2.25 ± 1.14 min). Complex formation with RafRBD was confirmed biochemically by a pulldown assay (Fig. S2 F, G), before and after concentration of the G-domain to the PM. Furthermore, PM concentration triggered increased phosphorylation of the canonical ERK and Akt downstream kinases, confirmed by Western Blot (Fig. S2 H-I). Together, these results show that the GTP-bound state of CD-GDom bypasses the requirement of upstream activation by GEFs or receptor tyrosine kinases, uncoupling the oncoprotein-mediated effects from upstream regulation and establishing the Opto-CID-KRASG12V system as a tool for acute, reversible and spatially controlled activation of KRASG12V signaling at the oncoprotein level.

### Acute, global activation of KRASG12V oncoprotein signaling reduces collective migration in MDCK monolayers

By using the Opto-CID-KRASG12V system, we could successfully activate KRASG12V oncoprotein signaling activity with spatial control; we next asked how the acute and global activation affects the collective cell behavior in MDCK cell monolayers. To address this question, we first tested whether CD-GDom can be concentrated at the PM simultaneously across all cells within the monolayer by incubating Opto-CID-KRASG12V MDCK cells with TMP-Cl, the uncaged version of the dimerizer (ChiroBlock, 1 µM), which recruits CD-GDom to the PM by binding to HaloTag and ecDHFR proteins upon cell entry without further activation. After incubation, PM concentration of CD-GDom reached a plateau at 90 min with slower kinetics (Fig. 2 A, B; t_1/2_ = 48 ± 21 min) compared to the light-induced Opto-CID system, consistent with the near-instantaneous uncaging of the dimerizer at the PM after light irradiation, bypassing the diffusion and equilibrium steps required for externally added small molecules to reach their intracellular targets. Subsequent addition of free TMP (10 µM) in excess reversed CD-GDom localization back to the cytoplasm within 5 min.

**Figure 2.**
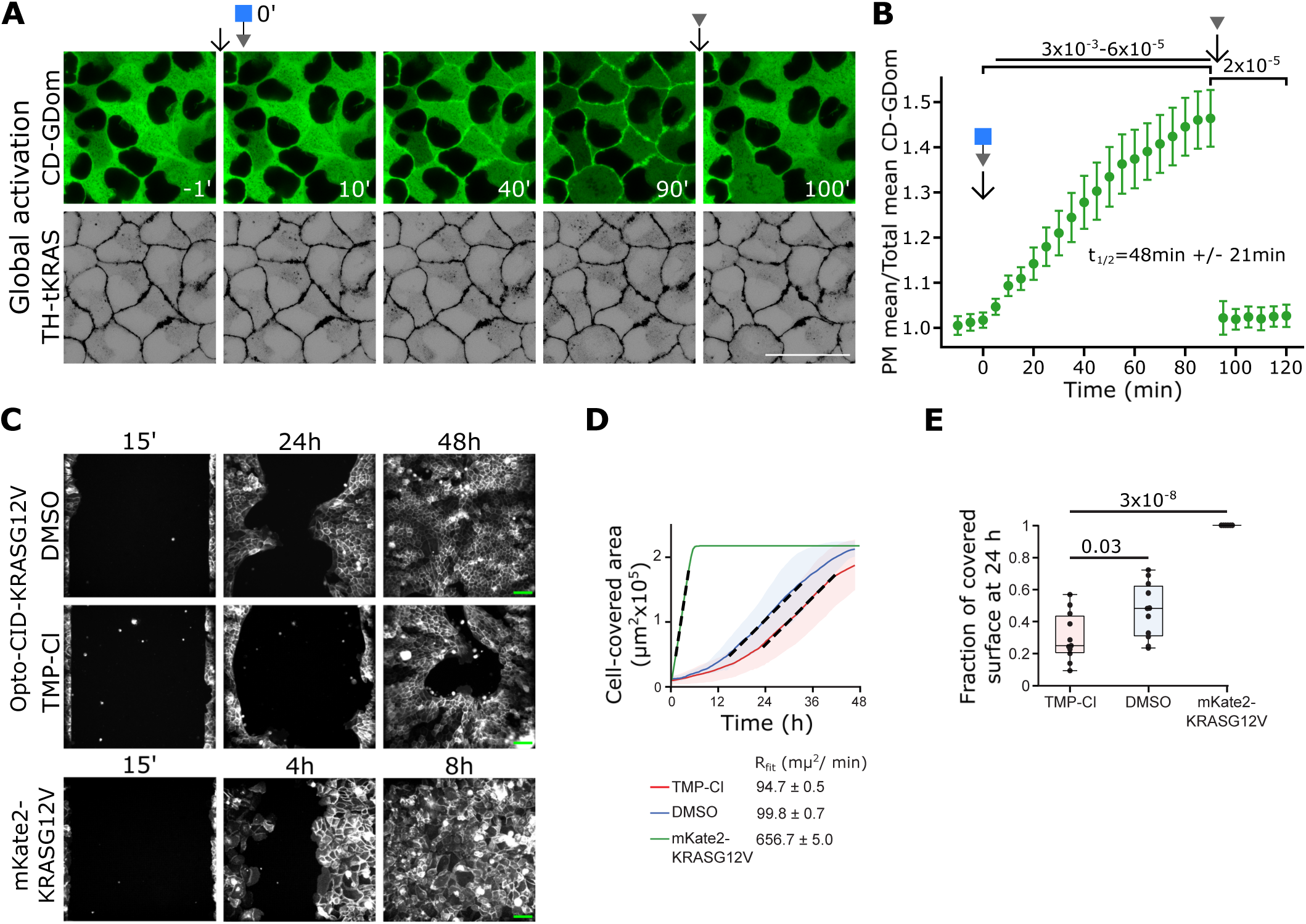
Acute, global CD-GDom PM concentration promotes decreased migration in MDCK cells. **(A)** Representative micrographs of CD-GDom (green) and TH-tKRAS (inverted grey) fluorescence at the indicated time points in Opto-CID-KRASG12V-MDCK cells before and after incubation with the TMP-Cl dimerizer (1 μM) and the subsequent TMP (10 μM) addition for reversibility after 10 min. **(B)** Scatter plot depicting the ratio of the CD-GDom mean intensity of PM over total (PM + cytoplasm) over time, showing the t_1/2_ time until reaching the maximum PM concentration. *p*-value was calculated by the Mann-Whitney U test with Bonferroni correction, with n = 15. **(C)** Representative snapshots of gap-closure assays (initial gap width 500 µm) in MDCK monolayers expressing the Opto-CID-KRASG12V construct treated with DMSO (Top) or TMP-Cl (middle), and mKate2-KRASG12V (bottom), acquired at 15 min, 24 h and 48 h after barrier removal (Opto-CID-KRASG12V) or at 15 min, 4 h and 8 h (mKate2-KRASG12V). **(D)** Time course of cell-covered area after barrier removal for TMP-Cl-treated (red), DMSO-treated (blue) and mKate2-KRASG12V expressing (green) monolayers, averaged over 11 positions (N = 6 wells) for TMP-Cl, 11 positions (N = 5) for DMSO and 6 positions (N = 4) for mKate2-KRASG12V. Linear regions were globally fit to obtain the slope R_fit (µm²/min), which quantifies the rate of gap closure. **(E)** Box plots of the fraction of the initial gap covered by cells 24 h after barrier removal for TMP-Cl, DMSO and mKate2-KRASG12V conditions, with individual positions shown as overlaid points. p-values were calculated using a two-sample t-test. Scale bar = 50 μm for **A** and 200 μm for **C**.

As oncogenic KRAS mutations leading to chronic oncoprotein activity have been shown to promote cellular migration in different cell types (Makrodouli *et al*, 2011; de Borja *et al*, 2025), we used a migration assay to investigate the collective migratory response to a global PM concentration of CD-GDom in MDCK cells, by adding either TMP-Cl or DMSO with barrier-removal between a confluent monolayer (Fig. 2 C; see Methods). The cells incubated with the dimerizer showed reduced migratory dynamics over 48 h with a significant decrease at 24 h compared to vehicle treatment (DMSO; Fig. 2 D, E), with no significant change in cell proliferation as determined by fraction of EdU+ cells (Fig. S3). In contrast, MDCK cells expressing a constitutively PM-localized KRASG12V oncoprotein (mKate2-KRASG12V) showed increased migratory behavior, closing the gap in less than 8 h (Fig. 2 C-E). This counterintuitive result can be explained by the acute and synchronous KRAS signaling within all the cells in the monolayer, overriding the spatial ERK activity waves shown to promote cell migration (Aoki *et al*, 2017) without cellular adaptations associated with chronic mutant KRAS expression. Furthermore, a previous study reported that the genomic KRASG12V mutation in MCF10A cells induces a dysregulation in population directional movement, leading to an overall decreased cell migration (Lee *et al*, 2021).

### Opto-CID-KRAS-G12V-mSIOs develop WT-like morphology

The reduced collective migration observed in MDCK cell monolayers showed the effect of acutely activating KRASG12V signaling at the oncoprotein level on a homogeneous cell population. However, in cancer, KRAS drives oncogenic transformation in 3D structures, where interaction between different cell types, including stem cell niches, shapes the response to oncoprotein signaling. To investigate the consequences of the acute effect of activated KRASG12V oncoprotein signaling in this context, we applied the Opto-CID-KRASG12V system to mouse small intestinal organoids (mSIOs), self-organizing 3D structures that recapitulate the organization of the intestinal epithelium (Sato *et al*, 2009).

Wild-type mouse small intestinal organoids (WT-mSIOs) consistently develop a crypt-villus morphological structure (Sato *et al*, 2009) with crypts containing the stem cell niche and the highly proliferative transit amplifying cells (CD44+; Wielenga *et al*, 1999) and the villus region containing mainly absorptive differentiated cells (Aldolase B+). While maintaining these distinctive regions, mSIOs morphology is highly heterogeneous, and oncoprotein activation has been shown to modify mSIOs morphology, mirroring intestinal tissue oncogenic transformation (Kotani *et al*, 2021; Soetje *et al*, 2026, preprint). To investigate the morphological consequences of acute KRASG12V oncoprotein signaling, we generated mSIOs stably expressing the Opto-CID-KRASG12V system (Opto-CID-KRASG12V-mSIOs). These organoids displayed CD-GDom and TH-tKRAS cytoplasm and PM distribution, respectively (Fig. 3 A), comparable to previous observations in MDCK cells (Fig. 1). Both chimeric proteins display heterogeneous distribution with a consistent fluorescence enrichment at the villus regions as compared to the crypts. This pattern may reflect differences in cell proliferation, protein turnover, or transcriptional regulation between the two tissue regions (George *et al*, 2008; Harnik *et al*, 2021).

**Figure 3.**
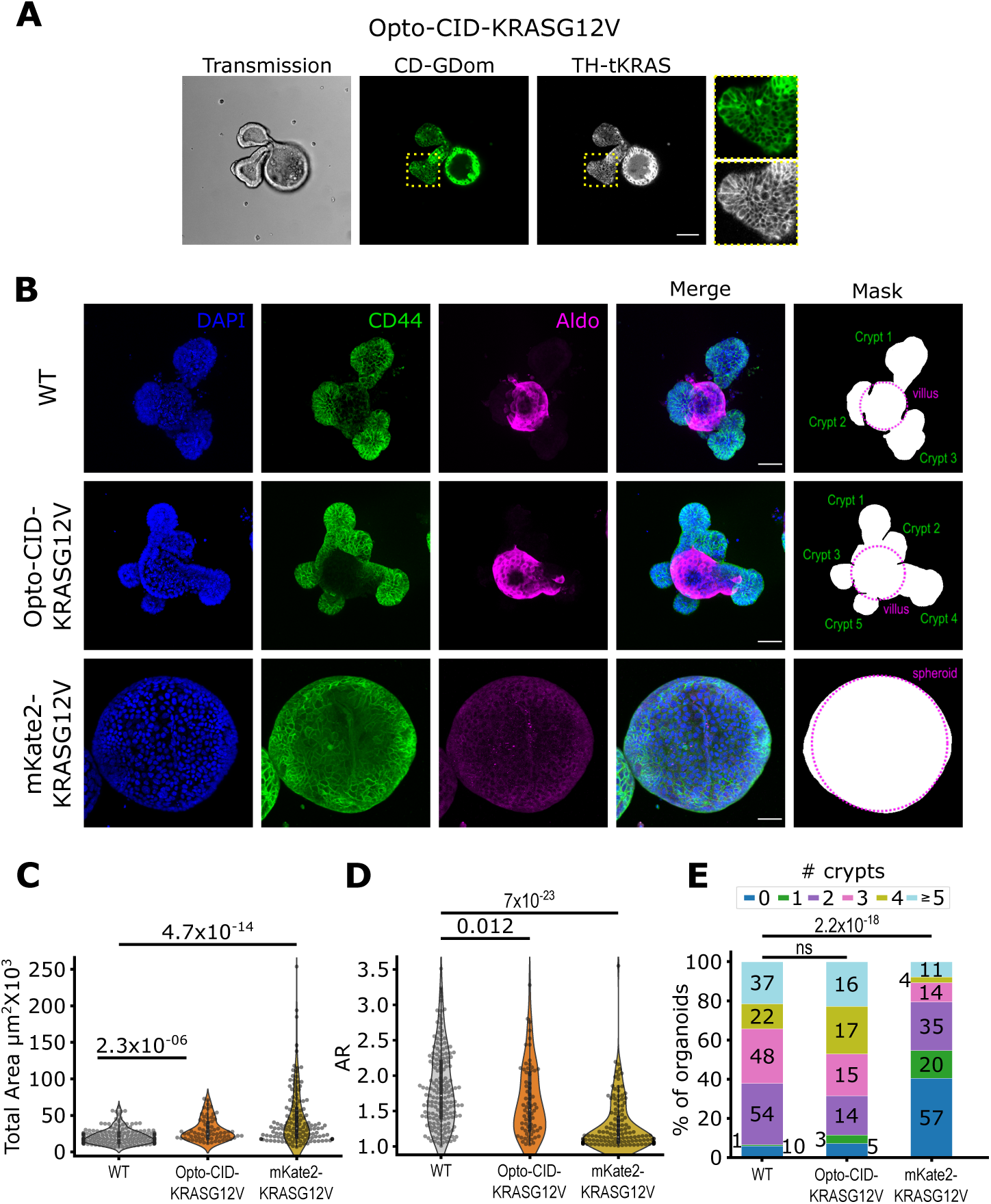
Opto-CID-KRASG12V-mSIOs develop a crypt-villus morphological structure similar to WT-mSIOs. **(A)** Representative CD-GDom (green), TH-tKRAS (inverted grey) fluorescence and Transmission micrographs from a 72 h developing mSIOs expressing the Opto-CID-KRASG12V DNA construct; magnifications of a crypt show the cytoplasmic and PM-tethered CD-GDom and TH-tKRAS, respectively. **(B)** Representative immunofluorescence images of WT-, Opto-CID-KRASG12V- and mKate2-KRASG12V-mSIOs depicting DAPI (blue), CD44 (green), Aldolase2 (Aldo; magenta) and the merge of them. **(B, right)** A mask was generated to segment the villus (magenta circumference) and the crypt regions for quantification; when no crypts were detected, mSIOs were considered as spheroids. **(C, D)** Violin plots depicting the morphological features: total Area and aspect ratio (AR) from WT, Opto-CID-KRASG12V, and mKate2-KRASG12V mSIOs. **(E)** Bar plot depicting the percentage of organoids with different numbers of crypts for all conditions; the numbers inside or next to the bars show the quantity of organoids with the corresponding number of crypts. *p*-value was calculated by the Mann-Whitney U test. n = 172, 70 and 141 for WT, Opto-CID-KRASG12V and mKate2-KRASG12V mSIOs, respectively. Scale bar = 50 μm.

We then tested whether our chimeric construct could affect the normal development of mSIOs, given that the cytoplasmic CD-GDom is constitutively GTP-bound, as shown in Fig. 1. For quantification, individual organoids were segmented with the villus region defined as the maximal inscribed circle and the crypts as protrusions outside this circle (Fig. 3 B, right; see Methods; Soetje *et al*, 2026, preprint); mSIOs lacking crypts were considered as spheroids. After 96 h incubation in standard enriched media (ENR), Opto-CID-KRasG12V-mSIOs develop a fully organized crypt-villus morphology with a slightly larger total area, decreased aspect ratio (AR), and a similar number of crypts compared to WT-mSIOs (Fig. 3 B-E). In contrast, about 40% of mSIOs expressing the constitutively PM-localized KRASG12V oncoprotein (mKate2-KRASG12V; Fig. S4) developed as spheroids showing disorganized and widespread expression of CD44 and no clear expression of differentiation marker Aldolase B (Aldo; Fig. 3 B). These mSIOs had an AR close to 1 and an increased total area, indicating that chronic KRASG12V oncoprotein signaling promotes aberrant tissue development, consistent with an increased proliferation reported for genomic KRASG12D mutation (Kotani *et al*, 2021). Altogether, Opto-CID-KRasG12V-mSIOs feature minor differences with non-transfected WT-mSIOs, likely explained by the GTP-bound state of CD-GDom, but without the loss of crypt-villus morphology observed under chronic expression of KRASG12V. Thus, Opto-CID-KRasG12V-mSIOs are suitable to study the consequences of acute KRASG12V oncoprotein signaling activation in organoid morphology.

### Acute and global KRASG12V signaling drives reversible crypt expansion in EGF-deprived mSIOs

To test whether PM concentration of CD-GDom affects mSIOs development, we globally induced KRASG12V oncoprotein signaling in Opto-CID-KRasG12V-mSIOs by incubation with TMP-Cl (1 μM, Fig. 4 A), leading to a gradual accumulation and reaching significant PM concentration of CD-GDom by 240-270 min (4-4.5 h) after incubation (Fig. 4 B). Reversibility was induced with the addition of the TMP molecule in excess (10 μM) leading to a rapid decline in PM localization in less than 30 min. As morphometric quantification required mature 96 h mSIOs, we confirmed that PM localization, and hence sustained KRASG12V signaling, was sustained during this developmental window by replacing the dimerizer at 48 h of incubation (Fig. S5 A, B).

**Figure 4.**
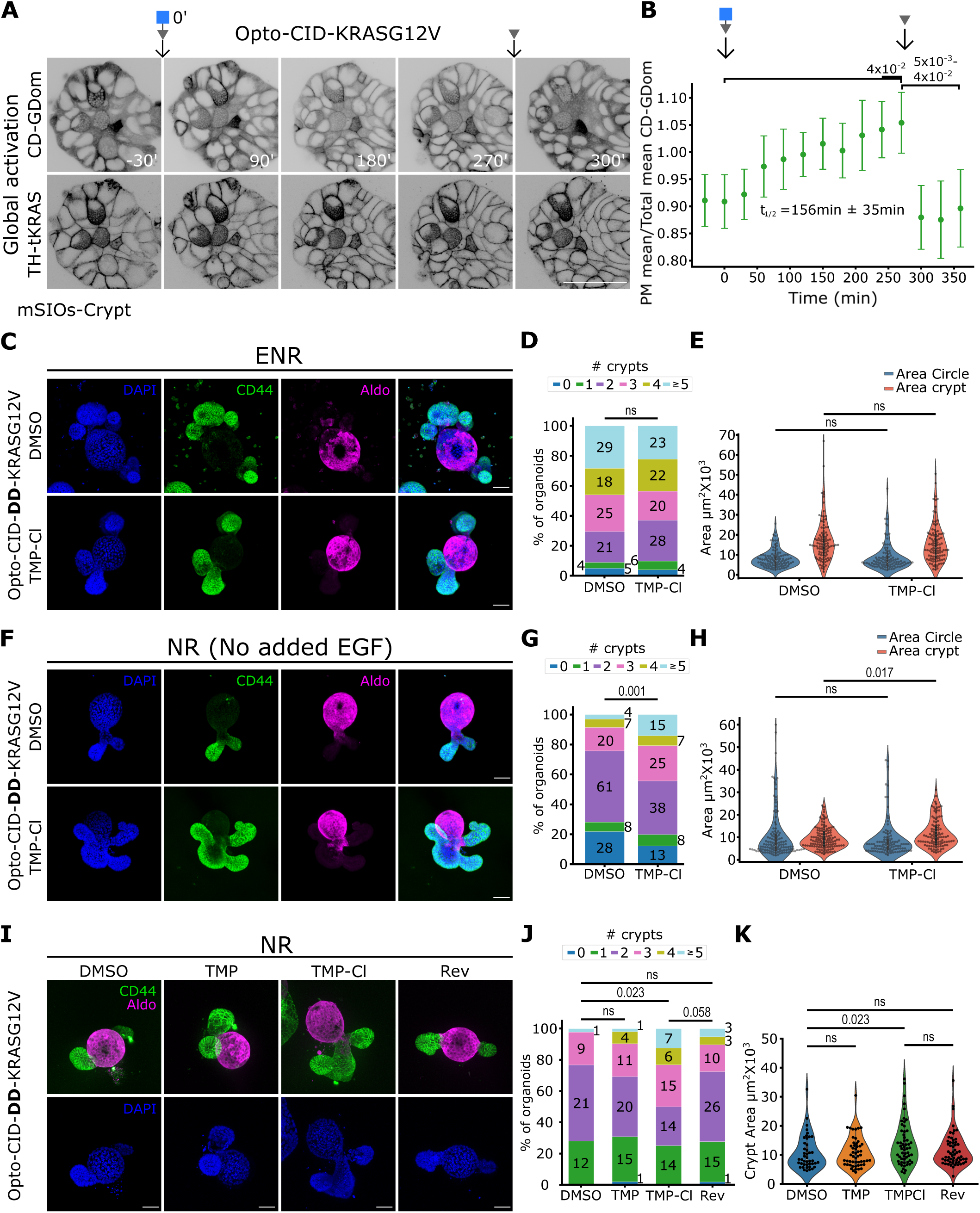
Global stabilization and PM concentration of DD-CD-GDom promote an increased crypt number and area in Opto-CID-DD-KRASG12V-mSIOs in EGF-deprived media. **(A)** Representative micrographs at the indicated time points from an Opto-CID-KRASG12V-mSIO crypt depicting CD-GDom (top) and TH-tKRAS (bottom) before and after TMP-Cl incubation (1 μM); TMP molecule (10 μM) was added after 270 min for reversibility of CD-GDom to the cytoplasm. **(B)** Scatter plot depicting the ratio of the CD-GDom mean intensity of PM over Total mean intensity (PM + cytoplasm) over time. **(C, F)** Representative immunofluorescence micrographs from Opto-CID-DD-KRASG12V mSIOs incubated for 96 h (media change at 48 h) with TMP-Cl or DMSO in normal EGF (EGF 50 ng/ml added; **C**), or EGF starvation media (NR; **F**); depicting DAPI (blue), CD44 (green), Aldolase2 (Aldo; magenta), and the merge of them. **(D, G, J)** Bar plot depicting the percentage of organoids with different numbers of crypts for all conditions; the numbers inside or next to the bars show the quantity of organoids with the corresponding number of crypts. Violin plots depicting the area corresponding to the villus region (Area Circle, **E, H**) and crypts **(E, H, K)**. Rev refers to Opto-CID-DD-KRASG12V mSIOs incubated for 48 h with TMP-Cl (1 μM), washed with TMP (10 μM), and incubated with DMSO for 48 h. *p*-value was calculated using the Mann-Whitney U test. n = (102, 103) and (128, 106) for (DMSO, TMP-Cl) incubated in ENR and NR media, respectively; n = 43, 52, 56 and 58 for DMSO, TMP, TMP-Cl and Rev treatments in **J** and **K**. Scale bar = 50 μm.

Opto-CID-KRasG12V-mSIOs grown in ENR media and treated with TMP-Cl exhibited no significant differences in size or number of crypts, compared to organoids growing in the same conditions without the dimerizer (Fig. S5 C-E), despite PM recruitment being sufficient to drive signaling (Fig. 1; Fig. S2) and migration defects (Fig. 2) in MDCK monolayers. We reasoned that two features of the mSIOs context could mask the phenotype. First, ENR media contains saturating EGF (50 ng/ml), which activates EGFR-driven KRAS signaling and have synergistic effects with mutant isoforms (Jiang *et al*, 2011; Hood *et al*, 2019; Ponsioen *et al*, 2021; Yokoi *et al*, 2025); saturating EGF also promotes endosomal degradation of EGFR in intestinal organoids (Boretto *et al*, 2024; Caracci *et al*, 2026, preprint), potentially limiting PM components necessary for KRAS signaling. However, Opto-CID-KRasG12V-mSIOs grown in NR media (no added EGF) developed smaller organoids, but without difference between TMP-Cl and DMSO vehicle controls (Fig. S5 F-H), indicating that EGF saturation alone does not account for the absence of phenotype differences. Second, the GTP-bound cytoplasmic CD-GDom could promote minor background signaling activity (Fig. 3 B-E) through complex formation with effectors (Fig. 1D, E). To decrease it, we introduced the ecDHFR destabilization domain (DD-ecDHFR; Iwamoto *et al*, 2010) into CD-GDom (Opto-CID-DD-KRASG12V; Fig. S5 I), driving its proteolytic degradation in the absence of TMP. TMP-Cl (1 μM) stabilized DD-CD-GDom in the villus region (similar to the stable ecDHFR version of the construct; Fig. S6 B, C, top) but not in the crypts. In contrast, incubation with TMP (1 μM), which stabilizes the DD-GDom without PM concentration, produced a significant increase in both regions (Fig. S6 B, top). This pattern may reflect downregulation of Opto-CID-DD-KRASG12V expression in crypts in response to PM-concentrated DD-CD-GDom (Fig. S5 I, right), leading to KRASG12V oncoprotein signaling, although other mechanisms cannot be excluded. For morphological analysis, we biochemically confirmed by Western Blot that DD-CD-GDom stabilization was sustained over 96 h by replacing media with TMP or TMP-Cl at 48 h (Fig. S5 J).

Opto-CID-DD-KRASG12V-mSIOs incubated with TMP-Cl in ENR media for 96 h showed no morphological difference compared to controls (Fig. 4 C-E). In contrast, when incubated in NR media, TMP-Cl-treated mSIOs developed more and larger crypts, decreasing the number of spheroids (Fig. 4 F-H). These results suggest that acute KRASG12V oncoprotein signaling activity, when both the EGF ligand promoted MAPK signaling and the cytoplasmic background activity of CD-GDom are reduced, leads to crypt morphological changes in a 96 h period. Furthermore, the increased number of crypts, also observed with endogenous KRAS genomic locus mutations (Kotani *et al*, 2021) could be explained by enhanced crypt fission in the mouse intestinal tissue as shown for the KRASG12D mutation (Snippert *et al*, 2014).

Finally, we tested whether the morphological changes observed are reversible upon removal of CD-GDom from the PM. We incubated Opto-CID-DD-KRasG12V-mSIOs for 48 h in NR media with TMP-Cl, then washed three times with TMP (10 μM) and re-cultured in NR media with the DMSO vehicle (Rev; Fig. 4 I-K). TMP-Cl-treated mSIOs consistently developed with more crypts and increased crypt area compared to controls (Fig. 4 I-K). Rev mSIOs showed a decreased number of crypts with respect to TMP-Cl-treated ones, with no significant differences from DMSO vehicle-treated mSIOs. As an additional control, Opto-CID-DD-KRasG12V-mSIOs incubated with the free TMP molecule (1 μM) showed no morphological difference compared to DMSO and Rev mSIOs, indicating that the phenotype depends specifically on PM concentration of CD-GDom and not on its stabilization in the cytoplasm. These findings suggest that the crypt morphological changes depend on sustained KRASG12V oncoprotein signaling, which is consistent with previous studies demonstrating that continuous KRAS activity is necessary for the maintenance of oncogenic transformation in different tissue contexts (Brummelkamp *et al*, 2002; Singh *et al*, 2009).

### Acute, local KRASG12V oncoprotein signaling in the developing crypt is sufficient to promote crypt morphology changes

As global oncoprotein signaling activation doesn’t represent the localized initial transformation leading to tumor formation (Jonkers & Berns, 2002), we next tested whether the spatial control of CD-GDom PM concentration in a defined subset of cells within otherwise unperturbed mSIOs is sufficient to alter its development, recapitulating in vivo focal tumor initiation.

To demonstrate the CD-GDom concentration to the PM after light irradiation in mSIOs, we pre-incubated Opto-CID-KRasG12V-mSIOs with NvocTMP-Cl in ENR media (1 μM, 16 ± 2 h), photoactivating a single developing crypt region (Fig. 5 A, orange dashed rectangle). Significant PM concentration of CD-GDom was observed by 2 h post-PA (Fig. 5 A, B), substantially slower than the second-time scale observed for MDCK cell monolayers (Fig. 1B, C). This difference likely reflects the technical challenges inherent to 3D structures like mSIOs, where densely packed small cells within a crypt, heterogeneous cell population, limited focal plane imaging, and reduced discrimination between PM and cytoplasmic fluorescence signals make the quantification of PM concentration difficult. Subsequent addition of TMP (10 μM) rapidly reverted CD-GDom enrichment to the cytoplasm within 1 h.

**Figure 5.**
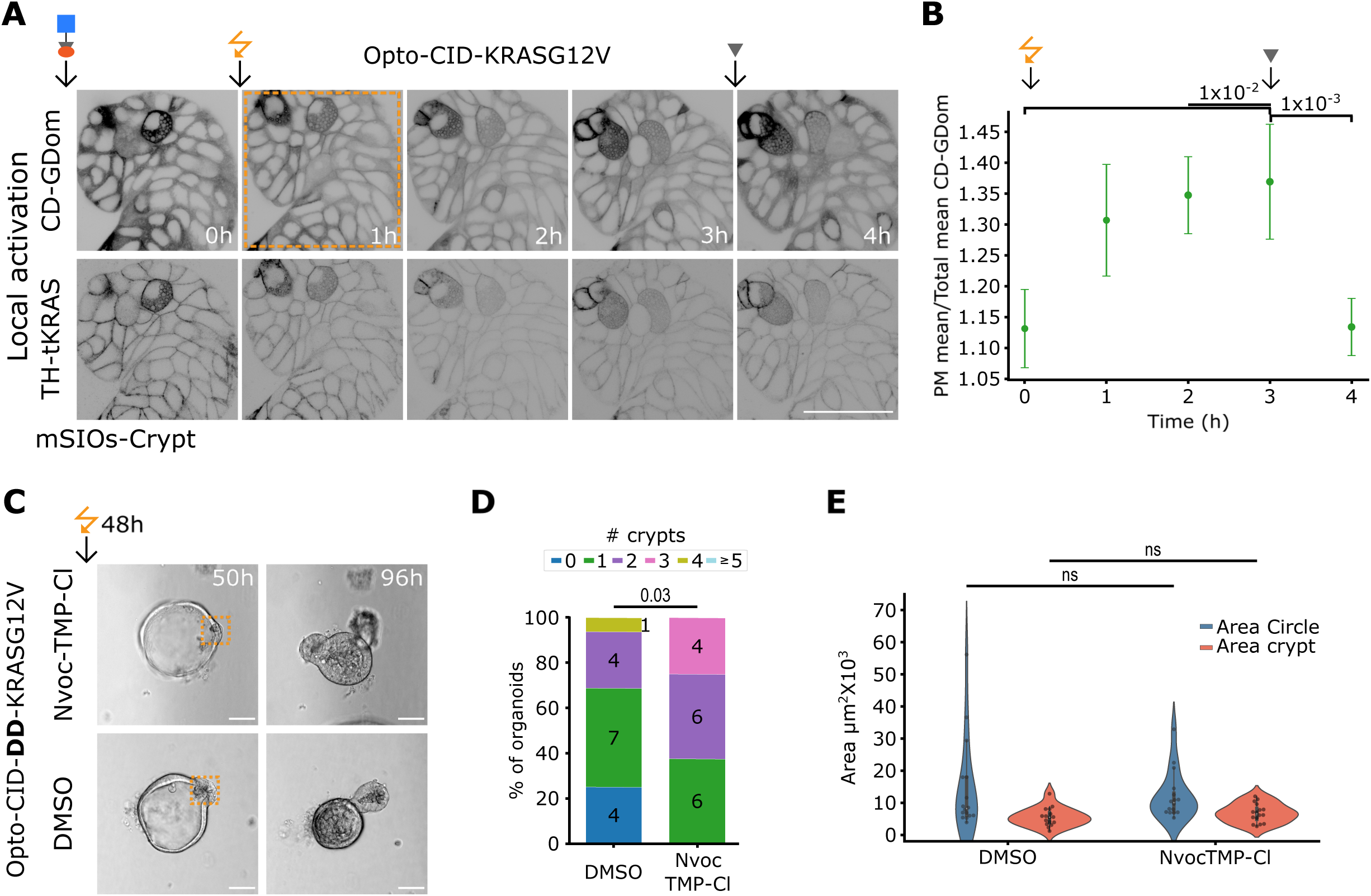
Local CD-GDom in a budding crypt promotes crypt-villus structure development in mSIOs in EGF-deprived media. **(A)** Representative micrographs at the indicated time points from an Opto-CID-KRASG12V-mSIO crypt depicting CD-GDom (top) and TH-tKRAS (bottom) after NvocTMP-Cl (1 μM) 16 ± 2 h incubation, PA via 405 nm light (within the complete field of view; orange dashed rectangle) at time 0 and the subsequent TMP (10 μM) addition for reversibility after 3 h. **(B)** Scatter plot depicting the ratio of the CD-GDom mean intensity of PM over Total mean intensity (PM + cytoplasm) over time. **(C)** Representative brightfield images of 48 h and 96 h Opto-CID-KRASG12V-mSIOs irradiated with 405 nm light at 48 h within the orange dashed rectangle. **(D)** Bar plot depicting the percentage of organoids with different numbers of crypts for all conditions; the numbers inside or next to the bars show the quantity of organoids with the corresponding number of crypts. **(E)** Violin plots depicting the area corresponding to the villus region (Area Circle) and crypts. *p*-value was calculated by the Mann-Whitney U test for **B** and **E**, with Bonferroni correction for B. n = 7 for **B** and 16 for DMSO- and NvocTMPCl-treated mSIOs for **E**. Scale bar = 50 μm.

To determine whether the localized KRASG12V oncoprotein signaling activation leads to morphological changes within the tissue, we incubated Opto-CID-DD-KRasG12V-mSIOs (reduced CD-GDom concentration) in NR media for 48 h (time until budding initiation), in order to photoactivate a single budding crypt without irradiating other regions within the mSIO (Fig. 5 C, dashed orange rectangle). mSIOs preincubated with Nvoc-TMP-Cl dimerizer consistently developed a crypt-villus morphological structure with a small but significant increase in the number of crypts as compared to DMSO vehicle controls (Fig. 5 D), in which some mSIOs developed as spheroids (∼25%). In contrast, crypt and villus areas showed no significant difference by local induction (Fig. 5 E). Despite technical challenges in the selectivity of light induction on 3D structures as mentioned above, our results suggest that local KRASG12V oncoprotein signaling activity within a single budding crypt, a stem cell niche, is sufficient to promote crypt formation across the mSIOs without affecting the overall size. This acts as a proof-of-principle of this technique; however, further advances in selective illumination of 3D tissues are necessary to strengthen our results or extend Opto-CID as a method to other 3D models.

## Discussion

Altogether, in this study we have developed a synthetic, two-component, opto-chemical system that decouples KRASG12V oncoprotein signaling from oncogene expression and experimentally implements the dimensionality reduction mechanism by which active KRAS recruits effector proteins to the PM. This system enables an acute, spatial, and reversible control of KRAS signaling in living tissues such as MDCK cell monolayers, where the Opto-CID-KRASG12V system alters ERK phosphorylation and dynamics, decreasing collective cell migration. In mSIOs, we added an additional layer of regulation by incorporating a destabilization domain into the GTP-bound CD-GDom that otherwise could lead to constitutive effector activation. In EGF-deprived conditions, CD-GDom PM concentration drove reversible crypt expansion, while local activation in a single budding crypt was sufficient to alter organoid development toward a crypt-villus morphology.

Synthetic biology tools based on split-protein approaches and/or CID have been extensively used to evaluate protein-protein interactions (Bae *et al*, 2024). Split-small CID GTPase system controlled KRAS, Cdc42, and RhoA signaling (He *et al*, 2024, 2025). More recently, the same group published a preprint showing spatial activation of Cdc42, RhoA, and Rac1 GTPases in single cells (Faulkner *et al*, 2025, preprint). Furthermore, light-inducible oligomerization has been successfully used to activate Wnt downstream signaling (Kaur *et al*, 2017; Lee *et al*, 2023; Repina *et al*, 2023; Soetje *et al*, 2026, preprint) and EGFR signaling (Suh et al, 2025). To our knowledge, our CD-GDom and TH-tKRAS are the first split-protein synthetic cell system reconstituted in living cells and organoids for reversibly and spatially controlling KRAS oncoprotein signaling.

Our opto-chemical approach is applicable to other RAS mutations, and to any oncoprotein that recruits effectors to the PM. It can also be employed in cells or organoids carrying tumor suppressor loss-of-function or oncogene gain-of-function mutations, enabling the study of oncoprotein contributions to transformation in a multiple mutation context. This is particularly relevant given that KRAS activation alone has been reported to produce modest or non-significant morphological changes in otherwise non-mutant tissue (Onuma *et al*, 2013; Brandt *et al*, 2019), consistent with the significant but modest changes in organoid morphology shown here (Fig. 4 and Fig. 5). Stronger transformation has been shown in combination with other mutations such as APC and p53 (Boutin *et al*, 2017; Onuma *et al*, 2013). Combination with complementary inducible systems for other oncoproteins, such as the opto-chemical oncomimetic β-catenin activation system, recently developed in our lab (Soetje *et al*, 2026, preprint) would further enable the study of the consequences of multiple acute oncoprotein signaling during tissue transformation.

Chimeric systems that enable acute, reversible, and spatially controlled activation of oncoprotein signaling activity in 3D structures across heterogeneous cell populations allow for a better understanding of an individual oncoprotein contribution to cancer development compared to conventional genomic mutation models. In these models, it is technically complicated to disentangle the immediate signaling consequences from the cellular adaptations of genomic instability that accumulate during sustained oncogene expression. Thus, continued advances in synthetic biology, light induction, and live imaging are essential to further understand the immediate signaling contribution of oncoproteins to cancer onset and development.

## Materials and Methods

### Plasmids and cloning

For the generation of MDCK cells and mSIOs stable lines, the cDNAs of interest were cloned into a *piggyBac-CAG-IRES-PuroR* vector (Wang *et al*, 2008) using the Gibson Assembly Master Mix (NEB, cat. # E2611s) following the manufacturer’s protocol. Briefly, vector and insert fragments were amplified by PCR using the Q5 High Fidelity DNA polymerase (NEB, cat. # M0491S) with up to 30 pb overlapping overhangs and assembled at 72 °C. DNA oligos for cloning were obtained from Thermo Fisher Scientific (RRID:SCR_008452). After assembly, all the DNA constructs were sequence-verified by Sanger sequencing (Microsynth Seqlab GmbH). The Opto-CID-KRASG12V construct (5.007 kb) is a combination of two fragments separated by the P2A-T2A ribosomal skipping sequence (Liu *et al*, 2017), consisting of CD-GDom and TH-tKRAS as explained in the text. For a destabilized CD-GDom, the ecDHFR N18T/A19V destabilization domain (DD-ecDHFR; Iwamoto *et al*, 2010) was used in substitution of ecDHFR. The booster-ERK FRET sensor, pCAGGS-4511NES was a gift from Michiyuki Matsuda (Addgene plasmid # 138375; http://n2t.net/addgene:138375; RRID:Addgene_138375) and was inserted between the two fragments of the Opto-CID-KRASG12V construct separated by a P2AT2A sequence. The CID-KRASG12V construct (3.765 kb), presented as supplementary material, is a combination of two fragments separated by the P2A ribosomal skipping sequence consisting of: 1) TF-GDom: Fused sequences of mTFP, a tandem of two FKBP37-V (Clackson *et al*, 1998a; Liu *et al*, 2014) and KRASΔHVR; and 2) BD-tKRAS: TagBFP, 2XecDHFR and tKRAS. Vector maps were designed and visualized using the SnapGene software (www.snapgene.com).

### Reagents

TMP: Trimethoprim (T7883; Sigma-Aldrich); SLF’-TMP: custom synthesis by ChiroBlockGmbH (Bitterfeld-Wolfen, Germany), chemical name: [(1R)-1-[3-[2-[3-[2-[2-[3-[[2-[4-[(2,4-diaminopyrimidin-5-yl)methyl]-2,6-dimethoxy-phenoxy]acetyl]amino]propoxy]ethoxy]ethoxy]propylamino]-2-oxo-ethoxy] phenyl]-3-(3,4-dimethoxyphenyl)propyl] (2S)-1-[2-(3,4,5-trimethoxyphenyl)butanoyl]piperidine-2-carboxylate; C63H85N7O17, M.W.= 1212.42 g/mol; NvocTMP-Cl: custom synthesis by ChiroBlock GmbH (Bitterfeld-Wolfen, Germany), chemical name: (4,5-dimethoxy-2-nitro-phenyl)methyl N-[4-amino-5-[[4-[2-[2-[2-[2-[2-(6-chlorohexoxy)-ethoxy]ethoxy]ethoxy]ethoxy]ethoxy]-3,5-dimethoxy-phenyl]methyl]pyrimidin-2-yl]carbamate; C39H56ClN5O14; M.W.=854.36 g/mol; TMP-Cl: custom synthesis by ChiroBlock GmbH (Bitterfeld-Wolfen, Germany), chemical name: N’-[2-[2-(6-chlorohexoxy)ethoxy]ethyl]-N-[3-[4-[(2,4-diaminopyrimidin-5-yl)methyl]-2,6-dimethoxy-phenoxy]-propyl]butanediamide; C30H47ClN6O7; M.W.= 639.20 g/mol.

### Cell culture and transfection

Madin-Darby canine kidney (MDCK) cells, (MDCK [NBL-2], ATCC® Number: CCL-34TM cells) were cultured in Dulbecco’s modified Eagle’s medium (DMEM; PAN-Biotech GmbH, Aidenbach, Germany) supplemented with 10 % fetal bovine serum (FBS; PAN-Biotech GmbH), 2 mM L-Glutamine and 1 % nonessential amino acids (NEAA, PAN-Biotech GmbH), and grown at 37 °C, 5 % CO_2_ in humidified incubators. The cells were transiently transfected with the Lipofectamine 3000 transfection reagent (Invitrogen, cat. #L3000001) following the manufacturer’s protocol, changing media 24 h after transfection. For stable cell line generation, a *piggyBac* vector (Wang *et al*, 2008) with the cDNA of interest and the *piggyBac* Transposase were co-transfected following the same protocol and selected with Puromycin (3 µg/ml) 24 h after transfection.

### Mouse Intestinal Organoid culture and transfection

Mouse Intestinal Organoids (mSIOs) were obtained from STEMCELL Technologies (STEMCELL Technologies, cat. # 70931), cultured in enriched media (ENR) containing: 1) advanced DMEM/F12 medium (AdDF, Thermo Fisher Scientific, Cat. # 12634010) supplemented with Hepes (Thermo Fisher Scientific, cat # 15630056), Glutamax (Thermo Fisher Scientific, cat # 35050038) and penicillin/streptomycin (Life Technologies, cat. # 15140122) (AdDF+++); and enriched with: 2) B27 (Life Technologies, cat # 17504044), 1.25 mM N-acetylcysteine (Sigma Aldrich, cat. # A9165 – 5g), 100 ng/mL Noggin (Peprotech, cat. # 250-38), 500 ng/mL R-Spondin 1 (Peprotech, cat. # 315-32) and 50 ng/mL of mouse epidermal growth factor (mEGF; Peprotech, cat # 315-09). NR media was generated without mEGF addition. For splitting, the organoids embedded in Matrigel (BD Corning, cat. # 356231) were resuspended in cold AdDF+++ medium, centrifuged at 4 °C and 80 g, disintegrated to crypts by pipetting up and down 50-80 times, centrifuged at 400 g, seeded in 30 μl Matrigel droplets into 24 polystyrene well plates (Sarstedt, cat. # 83.3922.500) and grown in ENR at 37 °C, 5 % CO_2_ in humidified incubators.

mSIOs were transfected by electroporation using the NEPA21 electroporator (Nepagene, Japan) as previously described (Fujii *et al*, 2015). In brief, mSIOs were seeded in 6 wells of a 24 polystyrene well plate and incubated with ENR, supplemented with CHIR99021 (Sigma-Aldrich) and Y-27632 (Hölzel Biotech, cat # M1817-50mg) (ENR++) for 24 h, adding 1.25 % DMSO (ENR+++) for the next 24 h. The day of transfection, mSIOs were dissociated to single cells by incubating them in TrypLE (Gibco, cat. # 12605010) supplemented with Y-27632 at 37 °C for 15 min, centrifuged at 4 °C and 400 g, washed twice with Opti-MEM (Gibco, cat. # 31985070), centrifuged and the resuspended with 100 μl of Opti-MEM containing the *piggyBac* vector with the cDNA of interest (10 μg) and the *piggyBac* transposase vector (3 μg) into a 2 mm gap cuvette (Xceltis, EC-002S) for electroporation. Poring/transfer pulse conditions were used as previously described (Fujii *et al*, 2015): Voltage – 175/20 V, Pulse length – 5/50 ms, Pulse interval – 50/50 ms, Number of pulses – 2/5, Decay rate 10/40 %, Polarity +/+/-. After electroporation, mSIOs were incubated in ENR++ for 30 min at room temperature, centrifuged, and resuspended in 5-6 Matrigel droplets and incubated with ENR+++. After 24 h, media were replaced with ENR++ for another 48 h, followed by ENR for growth and maintenance.

### mSIOs Immunofluorescence

mSIOs were cultured from single crypts in ENR or NR media using 8- and 4-well Lab-Tek Chambered Coverglass (Sarstedt, cat # 94.6190.802) for 96 h, replacing media at 48 h with the specified treatment. The organoids were fixed by the addition of PFA 4 % for 40 min at room temperature, washed three times with IF-Buffer (DPBS, Tritonx100-0.2 %, Tween20-0.05 %), permeabilized for 1 h (DPBS, Triton X-100 0.5%), and blocked for 1 h (IF-Buffer, 3 % BSA) at room temperature. The samples were incubated overnight at 4 °C with primary antibodies: CD44 (1:500; BD Pharmigen™, 560568) and Aldolase (1:500, Abcam, ab153828), subsequently washed 3 times with IF-Buffer and incubated for 1 h at room temperature in the dark with secondary antibodies: Alexa Fluor anti-Rat 546 (Invitrogen, cat. # A-11081) and Alexa Fluor anti-Rabbit 647 (Invitrogen, cat. # A-31573). Incubation with DAPI (1:1000, Cell Signaling Technology, cat. # 4083s) for 15 min was used for nuclei staining. Images were obtained using confocal laser scanning microscopy.

### Confocal laser scanning microscopy (CLSM)

CLSM was performed using a TSC SP8 microscope (Leica Microsystems, Wetzlar, Germany) with an adapted temperature-controlled chamber set at 37 °C and 5% CO_2_ for live imaging experiments. Objectives: HC PL APO CS2 20x/0.75 DRY, HC PL APO CS2 63X/1.4 OIL; Pinhole: 1.0 Airy units; Lasers: 405 nm: DMOD Flexible, 470 nm to 670 nm: white light laser Kit WLL2 + Pulsepicker; Excitation / Emission wavelengths: DAPI (405 / 414-448 nm), mTFP (470 / 484-512 nm), mCitrine (514 / 531-566 nm), mKate2 (594 / 606-674 nm), detected with hybrid detectors.

### MDCK cells migration assay

MDCK cells were seeded the day before the experiment into a culture-insert 2 well ibiTreat (ibidi, cat. # 81176) placed inside a 2 well Lab-Tek Chambered Coverglass (Sarstedt, cat. # 94.6190.202) or a 24 well glass bottom plate (Cellvis, cat. # P24-1.5H-N), pre-coated with fibronectin (cat. # f0895-1mg). The day of the experiment, the cells were incubated with TMP-Cl (1 μM) or DMSO (1:10000) for 2 h in imaging media without phenol red (PAN-Biotech GmbH, cat. # P04-03588). Subsequently, the insert was removed to let the cells migrate for 48 h. Fluorescence images were taken every 10 min during the complete experiment, unless otherwise specified, using an Olympus IX81 inverted microscope (Olympus Life Science), with a pE-4000 LED illumination system (CoolLED), a Stage Top Incubator (ibidi) and ORCA-Quest qCMOS camera (Hamamatsu Photonics, Japan). To visualize proliferation, the cells were incubated at 37 °C, 5 % CO_2_ in humidified incubators without live microscopy for 20 h, as MCDK monolayers migrated faster without light irradiation. Proliferation was visualized by using the EdU staining assay (Click-iT EdU Alexa Fluor 647, Invitrogen, C10340), following manufacturer instructions and incubating the cells with EdU for 1 h prior to fixation.

### MDCK cells and mSIOs live imaging after chemical-global CD-GDom PM concentration

MDCK cells and mSIOs from single crypts were seeded in 8-well Lab-Tek Chambered Coverglass 48 h and 72 h prior to the experiment, respectively, unless otherwise specified. A TMP-Cl 5x stock solution (5 μM) was added in a 1:5 volume ratio to achieve a final concentration of 1 μM. For reversibility experiments, a 5x TMP stock solution (50 μM) was added in a 1:5 volume ratio to achieve a final concentration of 10 μM at the time stated in the experiment. Images were taken using CLSM.

### MDCK cells and mSIOs live imaging after Opto-chemical-local CD-GDom PM concentration

MDCK cells and mSIOs were seeded in a 24-well glass bottom plate (Cellvis, cat. # P24-1.5H-N). MDCK stable cell lines were seeded or transiently transfected 48 h prior to the experiment. mSIOs were seeded from single crypts 72 h prior to the quantification of PM CD-GDom concentration experiment and 48 h for morphometric analysis after photoactivation (PA) of a single budding crypt. MDCK cells and mSIOs were incubated inside the incubator at 37 °C with NvocTMP-Cl in DMEM complete growth media (10 μM, for 1 h) and in NR/ENR media (1 μM, overnight), respectively. Not bound dimerizer was removed by 3 times wash cycles of 1) 3x 1 min incubation with warm PBS or AdDF+++ solution followed by 2) 30 min incubation in CGM or NR/ENR media, for MDCK cells and mSIOs, respectively. For reversibility experiments, a 5x TMP stock solution (50 μM) was added in a 1:5 volume ratio to achieve a final concentration of 10 μM at the time stated in the experiment.

Photo-uncaging was performed using a TSC SP8 microscope, by illumination of a region of interest (ROI) with light at 405 nm wavelength with a 63x/1.4NA oil immersion objective. The illumination dose ≍ 6 J cm2 was used to achieve the maximal degree of dimerization as previously stated (Chen *et al*, 2018a), this was achieved by using a 405 nm laser at 125µW power (405@100%), scanning the ROI 3 times, with zoom 3 and with a speed of 100 Hz in a 512x512 image.

### Förster resonance energy transfer (FRET)

Booster-ERK contains an ERK substrate peptide that, upon phosphorylation by active ERK, binds to the proline-directed WW phospho-binding domain, triggering conformational rearrangements of the sensor that increase FRET. The nuclear export signal (NES) restricts the sensor to the cytoplasm, thereby enabling readout of cytoplasmic ERK activity. To include Booster-ERK in the Opto-CID-KRASG12V system, mCitrine and mTFP were reassigned to TH-tKRAS and CD-GDom, respectively, to minimize spectral overlap between cytoplasmic mCitrine and the mKOk fluorophores.

To examine the sensor response to KRASG12V plasma membrane recruitment, MDCK cells expressing Opto-CID-KRASG12V-Booster were incubated with NvocTMP-Cl or DMSO in starvation media (0.5% FBS), and mKOk donor excitation was used to record both mKOk and mKate2 emission for at least 30 minutes before and after photoactivation. The mCitrine channel was used for cell segmentation, enabling extraction of single-cell mKate2/mKOk emission ratios. CLSM Excitation / Emission wavelengths were as follows: mTFP (470 / 475-505 nm), mCitrine (514 / 525-535 nm), mKOk (550 / 560-570), mKate2 (550 / 650-755 nm).

### Immunoblotting and Pulldown assays

MDCK cells grown in 6-well-standard plates (SARSTEDT) and mSIOs derived from single crypts grown in 24-well-suspension plates (SARSTEDT) were lysed using RIPA cell lysis buffer (50 mM Tris–HCl pH 7.9, 150 mM NaCl, 1 % IGEPAL, 0.5 % Na-deoxycholate, 20 mM NEM) and modified RIPA lysis buffer (50 mM Tris HCl, pH 7.4, 150 mM NaCl, 1 mM EDTA, 1 mM EGTA, 0.5 % Triton X-100, and 0.5 % sodium deoxycholate), respectively. Both were supplemented with Complete Mini EDTA-free protease inhibitor (Roche Applied Science, Heidelberg, Germany) and phosphatase inhibitor cocktail 2 and 3 (1:100, P5726 and P0044, Sigma-Aldrich), shock frozen, and stored at -70 °C. mSIOs tissue was then dissociated using a 27G-gauge needle and a 1 mL syringe and centrifuged at 12,000 rpm for 10 min at 4 °C. Protein concentration was determined by BCA (Thermo Scientific, 23235); 20 μg of protein was loaded in 12% SDS/PAGE gels and transferred into PVDF membranes. Membranes were blocked using Intercept (TBS) blocking buffer (LI-COR BioScience, 927-60001) and incubated with primary antibodies overnight: PanRAS (1:1000; Invitrogen, Cat. # MA1-012), HaloTag (1:1000; Promega, cat # G9211), pERK (Cell Signaling Technology, cat. # 4370), pAkt (Cell Signaling Technology, cat. # 9271) and α-Tubulin (1:10000; Sigma Life Sciences, cat # T6074) were used. Secondary antibodies: IRDye 680RD and 800CW Donkey Anti-mouse IgG (LI-COR, cat # 926-68072 and 926-32212), IRDye 680RD and 800CW Donkey anti-Rabbit IgG (LI-COR, cat # 926-68073 and 926-32213).

For the pulldown assay, thawed cell lysate containing 200 µg of total protein per sample (with a minimum threshold of 100 µg when lysate availability was limited) was transferred to pre-chilled 1.5 mL microcentrifuge tubes containing 30 µL of purified 3x-RafRBD-coupled magnetic beads resuspended in cell lysis buffer. Samples were incubated for 30 min at 4 °C under continuous rotation. Total protein concentration was normalized across all samples within each experimental replicate. Nucleotide-loaded control samples pre-treated with either GDP (negative control) or GTPγS (positive control) were processed under identical conditions. Following incubation, each reaction was subjected to three sequential wash steps with 500 µL of cell lysis buffer, with complete removal of residual buffer after the final wash. For elution and downstream analysis by SDS-PAGE and western blotting, bound proteins were resuspended in 50 µL of 2X SDS sample buffer and denatured by heating at 95 °C for 10 min. Tubes were briefly cooled on ice and centrifuged for 10 seconds, after which the supernatant was collected using a magnetic separation stand and transferred to fresh pre-chilled 1.5 mL tubes for storage at -20 °C prior to SDS-PAGE. Throughout all wash and separation steps, a DynaMag-2 magnetic stand (Invitrogen) was employed to immobilize the magnetic beads against the tube wall, circumventing the need for centrifugal separation. Aspiration of supernatants was performed using a Vacusafe comfort vacuum pump in conjunction with a 200 µL pipette tip.

### Image analysis and quantification

To quantify the PM concentration of CD-GDom or TF-GDom in cells and mSIOs, a PM mask was generated for each time point from the selected cell population by applying a local intensity threshold to TH-tKRAS or BD-tKRAS. The fraction of GDom fluorescence signal at the PM was calculated as the ratio of the mean from GDom-positive pixels corresponding to the PM mask over the mean of total GDom-positive pixels. All the analyses were performed using Python 3.11 (python.org). Image processing and quantification were carried out with the following libraries: imageio (pypi.org/project/ImageIO/), NumPy (numpy.org), pandas (pandas.pydata.org), scikit-image (scikit-image.org), SciPy (pypi.org/project/scipy/). Data visualization was performed using matplotlib (matplotlib.org) and seaborn (seaborn.pydata.org).

Morphometric analysis of mSIOs was performed in accordance with the methodology previously described by our group (Soetje *et al*, 2026, preprint). In brief, micrographs of mSIOs acquired by confocal or widefield microscopy were processed in FIJI (fiji.sc). Individual organoid masks were generated by applying the “Li Dark” and “Otsu” automatic threshold algorithms to confocal and widefield micrographs, respectively, followed by manual curation to correct segmentation errors. Identification of the villus-like region within the mSIOs was achieved using the “Max. Inscribed Circle” function (Legland *et al*, 2016) with manual correction applied when needed. To define the presence of a crypt, we linearized the perimeter of the organoid by calculating the distance from the max. inscribed circle to each perimeter pixel. Crypts were detected as peaks on the linearized data using the “findpeaks” command in MATLAB. Specifying parameters: minimal peak prominence 15 μm; maximal peak width 1/3 of perimeter.

## Acknowledgements

This project was supported by the Max Planck Society for the Advancement of Science and ERC-AdG-322637 (SPATONC). We would like to thank Manuela Grygier and Jutta Luig for their assistance in cloning and plasmid preparation; Roger Goody, Jan Hübinger and Malte Schmick for critically reading the manuscript, feedback and discussions. Sven Müller and Michael Schulz for technical support for microscopy.

Prof. Dr. Philippe I.H. Bastiaens passed away on May 15, 2025. All authors agree that his inclusion as a co-author is appropriate to honor his intellectual contribution to this work.

## Author contributions

Luis Manuel Muñoz-Nava: Conceptualization and evolution of the project; experimentation in cells and organoids; methodology; investigation; data analysis; visualization; data curation; writing (original draft); reviews and editing.

Maria Bespalova: Experimentation in cells; methodology; investigation; data analysis; visualization; data curation; writing.

Mario O. Caracci: Experimentation in organoids; methodology; investigation; data analysis; data curation; writing.

Kitso Ata Kewagamang: Experimentation in cells; methodology; data analysis; visualization; data curation.

Birga Soetje: Investigation; methodology; data analysis.

Sabrina Seidler: Investigation.

Kirsten Michel: Investigation.

Holger Vogel: Conceptualization.

Janicke Maerz: Experimentation in organoids; data analysis.

Philippe I. H. Bastiaens: Conceptualization and evolution of the project; supervision; methodology; investigation; project administration; funding acquisition.

## Data Availability

Python and Matlab scripts can be provided on reasonable request.

