## Supplementary Material for "Reversible Opto-Chemical activation of KRASG12V signaling with near single-cell precision"

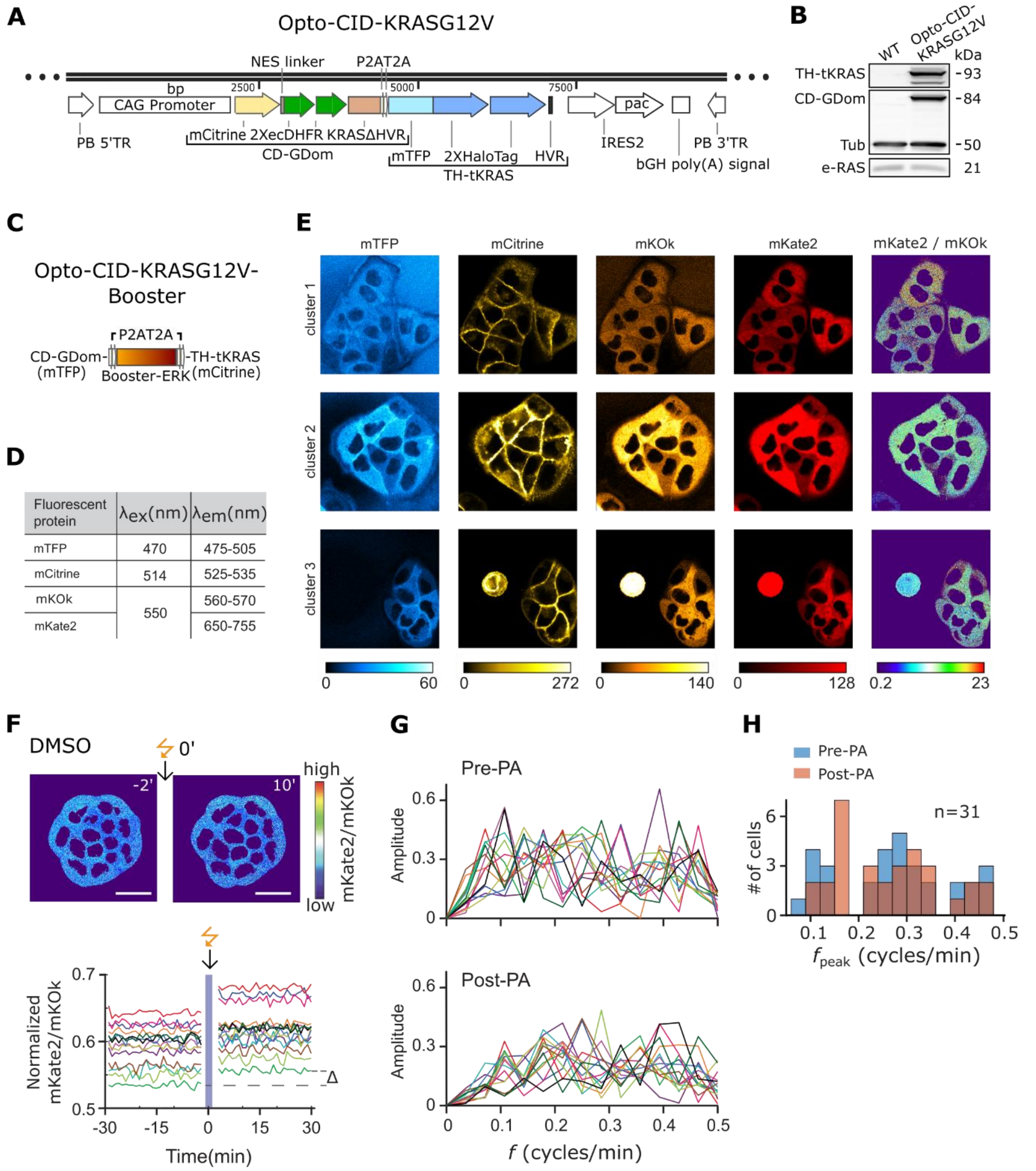


**Figure S1. Opto-CID-KRASG12V system DNA construct and the incorporation of the Booster-ERK FRET sensor.**

**(A)** Illustration depicting the fragment of the Polycistronic PiggyBac DNA vector encoding the Opto-CID-KRASG12V construct. bp: base pairs. From left to right: PB5’TR: piggyBac inverted terminal repeat; CAG: CMV-chickenβactin promoter sequence; CD-GDom (mCitrine-2XecDHFR-KRASG12VΔHVR); P2AT2A; TH-tKRAS (mTFP-2XHaloTag-tKRAS); IRES2: internal ribosomal entry site sequence for translation initiation; pac: Puromycin N-acetyltransferase expression cassette for selection; bGH poly(A) signal: bovine growth hormone polyadenylation signal termination sequence; PB3'TR: piggyBac inverted terminal repeat. **(B)** Western blot analysis from Opto-CID-KRASG12V-MDCK and MDCK WT cell lysates using antibodies against panRAS, HaloTag, and Tubulin. **(C)** Illustration showing the incorporation of the Booster-ERK FRET sensor to the Opto-CID-KRASG12V DNA construct (mCitrine and mTFP were reassigned to TH-tKRAS and CD-GDom, respectively, to minimize spectral overlap between cytoplasmic mCitrine and the mKOk fluorophores). **(D)** Excitation and emission wavelengths for all fluorescent proteins. **(E)** Confocal micrographs depicting mTFP, mCitrine, mkOƙ, mKate2, and the mKate2/mkOƙ ratio after NvocTMP-Cl incubation and 405 nm light irradiation for 3 different cell clusters. A color bar shows pixel intensity for each channel. **(F)** Top: Representative mKate2/mKOk images from Opto-CID-KRASG12V-Booster-MDCK cells incubated with DMSO, acquired 2 min before and 10 min after the 405-nm PA; scale bar 20 μm. Bottom: normalized single-cell time courses of the mKate2/mKOk; each trace was corrected for a global trend and rescaled to its first time point; the blue shaded area indicates PA, Δ denotes the PA-induced change in mKate2/mKOk, and $\Delta_{mean}$ indicates the mean change across all traces shown. **(G)** FFT amplitude spectra of the normalized mKate2/mKOk oscillations as a function of frequency, $f$, calculated from the pre-PA and post-PA segments of the time courses shown in **F**; colors were preserved between the time-course and FFT plots for each cell. **(H)** Distributions of peak frequencies, $f_{\text{peak}}$, derived from pre-PA (blue) and post-PA (orange) FFT spectra for DMSO-treated cells (n=31, from 3 independent cell clusters including cells shown in **F**). Each $f_{\text{peak}}$ represents the frequency of maximal FFT amplitude from an individual cell's spectrum.

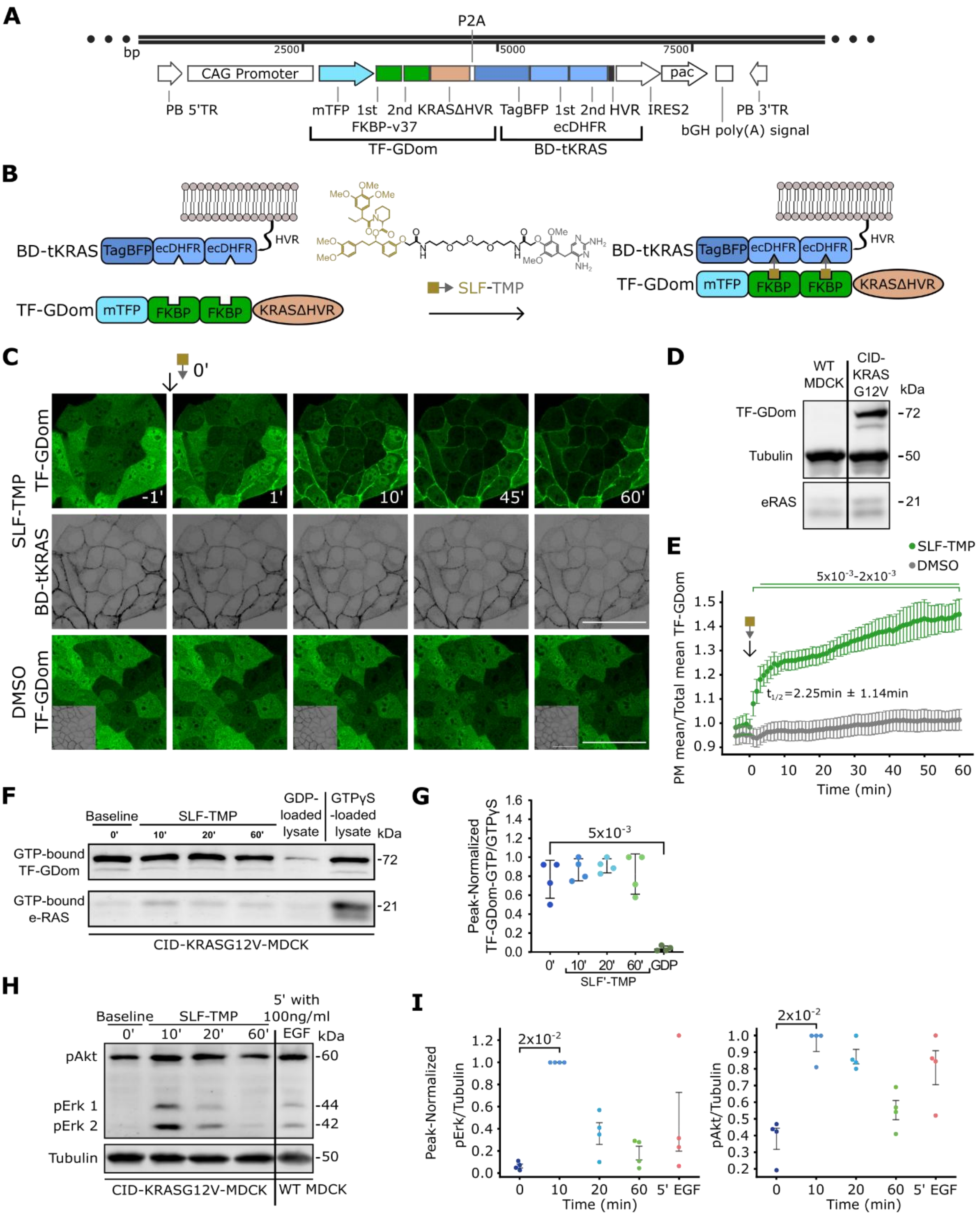


**Figure S2. Chemical-genetic orthogonal control of KRASG12V oncoprotein signaling activity by its concentration at the PM.**

**(A)** Polycistronic piggyBac DNA vector encoding the CID-KRASG12V construct. bp: base pairs. From left to right: PB5’TR: piggyBac inverted terminal repeat; CAG: CMV-chickenβactin promoter sequence; TF-GDom (mTFP-2XFKBP-KRASG12VΔHVR);

mTFP, FKBP: tandem of two FKBP36V molecules (a F36V mutant of FKBP, referred as FKBP) and KRASG12VΔHVR; P2A; BD-tKRAS (TagBFP-2XecDHFR-tKRAS); TagBFP: Tag-Blue fluorescent protein sequence; 2XecDHFR and tKRAS; IRES2: internal ribosomal entry site sequence for translation initiation; pac: Puromycin N-acetyltransferase expression cassette for selection; bGH poly(A) signal: bovine growth hormone polyadenylation signal termination sequence; PB3'TR: piggyBac inverted terminal repeat. **(B)** Scheme of the translated chimeric system depicting, from left to right: TF-GDom cytoplasmic and BD-tKRAS PM localization; SLF molecule (gold square) conjugated to the TMP molecule (gray triangle) that concentrates TF-GDom to the PM by binding to FKBP and ecDHFR molecules, respectively. **(C)** Representative micrographs from CID-KRASG12V-MDCK cells depicting TF-GDom and BD-tKRAS fluorescence at the indicated time points before and after SLF-TMP (2μM) or DMSO (1:10000) incubation. **(D)** Western blot analysis from CID-KRASG12V-MDCK and MDCK WT cell lysates using antibodies against panRAS and Tubulin. **(E)** Scatter plot depicting the ratio of the TF-GDom mean intensity of PM over Total mean intensity (PM + cytoplasm) with a $t_{1/2}$ = 2.25 min ± 1.14 min. n = 12 and 10 for SLF-TMP and DMSO, respectively. **(F, G)** Western blot analysis and quantification from CID-KRASG12V-MDCK cell lysates following a GST-3x-RafRBD pull-down without and with SLF-TMP (2 μM) incubation at the indicated time points using the panRAS antibody; GDP and GTPyS were used as negative and positive controls, respectively. **(H)** Western blot analysis and quantification from CID-KRASG12V-MDCK and MDCK WT cell lysates using antibodies against panRAS, pAkt, pERK, and Tubulin before and after SLF-TMP or EGF 100 ng/ml incubation. **(I)** Relative protein levels normalized to the highest value per experiment. *p*-value was calculated by the Mann-Whitney U test for all experiments and with Bonferroni correction for **E**. n = 4 for all Western Blot experiments. Scale bar = 50 μm.

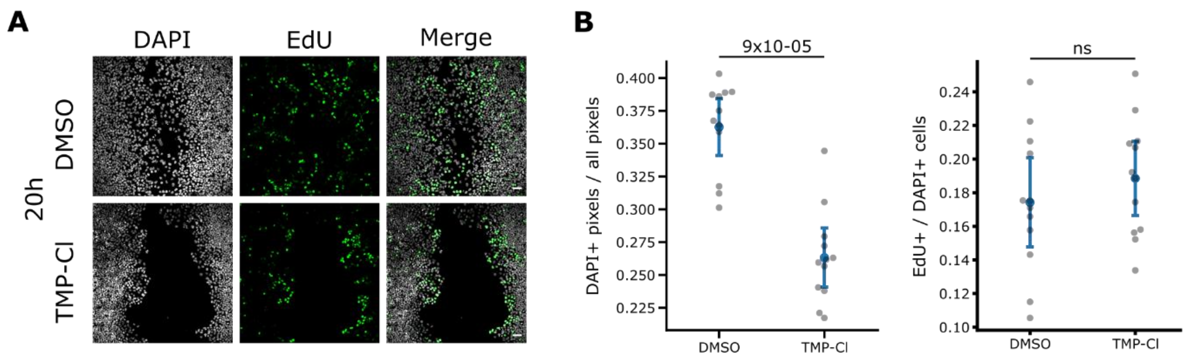


**Figure S3. Acute, global CD-GDom PM translocation does not affect cell proliferation.**

**(A)** Immunofluorescence images showing DAPI and EdU staining for Opto-CID-KRASG12V-MDCK cells after 20 h of migration incubated with DMSO or TMP-Cl. **(B)** Swarmplots showing DAPI+ pixels / all pixels within the field of view and the fraction of Edu+ pixels / DAPI pixels. *p*-value was calculated by the Mann-Whitney U test. n = 12. Scale bar = 50 $\mu$m.

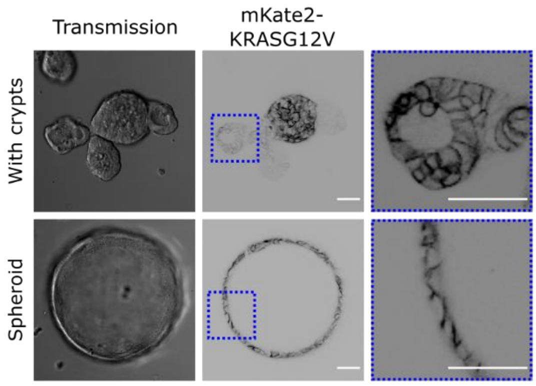


**Figure S4. Constitutive PM localization of mKate2-KRASG12V in mKate2-KRASG12V-mSIOs developed crypt-villus structures or spheroids.**

Representative micrographs from mKate2-KRASG12V-mSIOs at 72 h after single crypt seeding, depicting transmission and mKate2-KRASG12V fluorescence from a mSIO with and without crypts (spheroid), top and bottom, respectively. A magnification (blue dashed square) shows the accumulation of the oncoprotein to the PM (right).

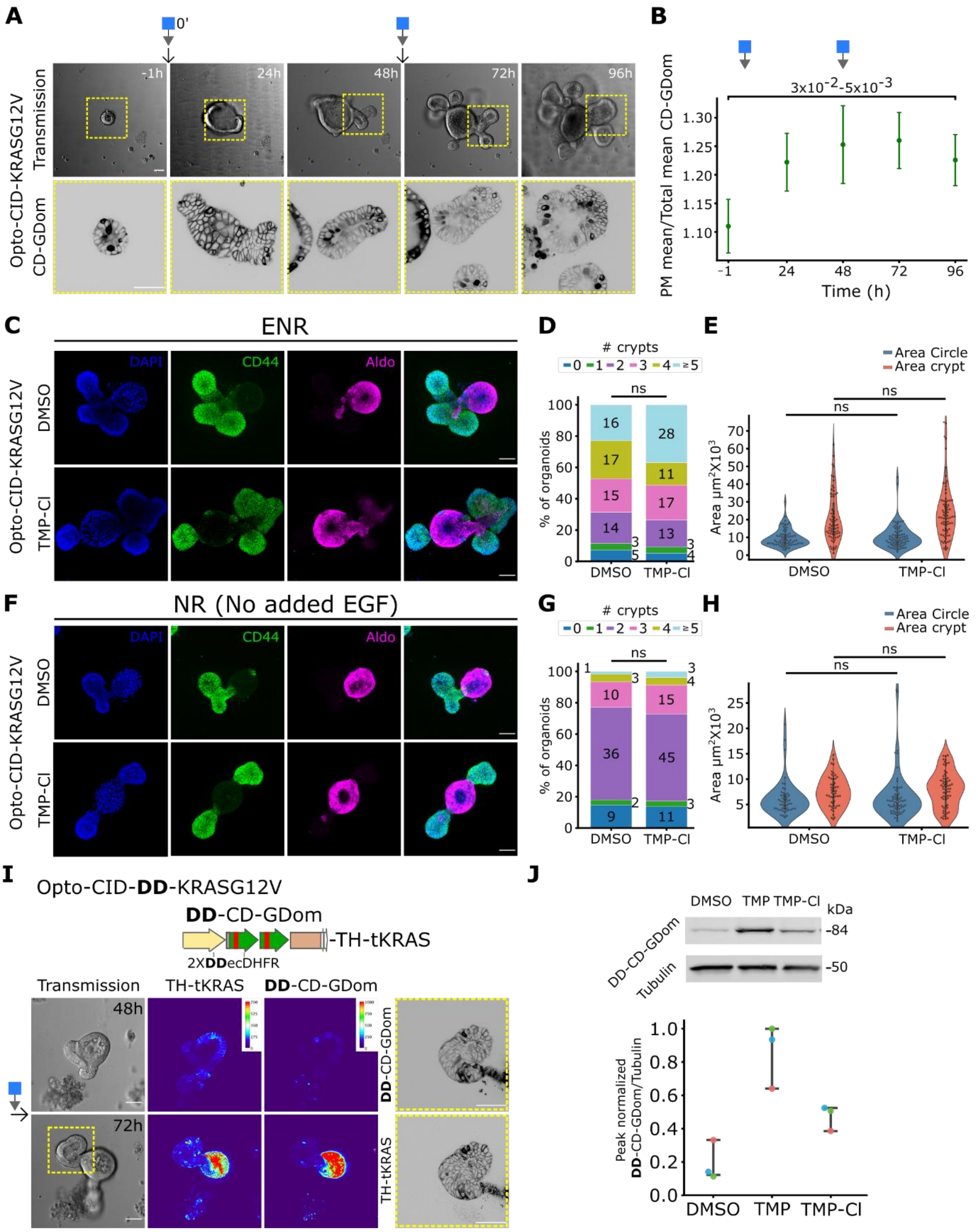


**Figure S5. Acute and global PM concentration of CD-GDom is not sufficient to induce morphological changes in Opto-CID-KRASG12V-mSIOs.**

**(A)** Representative transmission (top, 20X air objective) and CD-GDom (bottom, 63X oil immersion objective) micrographs at the indicated time point from an Opto-CID-

KRASG12V-mSIO before and after TMP-Cl incubation (1 μM, 0 and 48 h). **(B)** Scatter plot depicting the ratio of the CD-GDom mean intensity of PM over Total mean intensity (PM + cytoplasm) over time. **(C, F)** Representative immunofluorescence micrographs from Opto-CID-KRASG12V mSIOs incubated for 96 h (media change at 48 h) with DMSO or TMP-Cl in normal ENR (EGF 50 ng/ml added; **C**), or EGF starvation media (NR; **F**); depicting DAPI (blue), CD44 (green), Aldolase2 (Aldo; magenta), and the merge of them. **(D, G)** Bar plot depicting the percentage of organoids with different numbers of crypts for all conditions; the numbers inside or next to the bars show the quantity of organoids with the corresponding number of crypts. **(E, H)** Violin plots depicting the area corresponding to the villus region (Area Circle) and crypts. **(I, top)** Scheme depicting Opto-CID-DD-KRASG12V DNA construct bearing the DDecDHFR destabilization domain. **(I, bottom)** Representative transmission and heatmap TH-tKRAS, CD-GDom fluorescence micrographs of Opto-CID-DD-KRASG12V-mSIOs immediately before (48 h) and one day after (72 h) TMP-Cl (1 μM) addition. Showing a magnification (63X oil immersion objective, right) corresponding to one crypt (yellow dashed rectangle) 24 h after TMP-Cl addition. **(J)** Western Blot analysis and quantification from Opto-CID-DD-KRASG12V-mSIOs lysates incubated with DMSO, TMP, or TMP-Cl for 96 h with media change at 48 h, using an antibody against pan-Ras detecting DD-CD-GDom and against Tubulin. *p*-value was calculated by the Mann-Whitney U test. n = (70, 76) and (61, 81) for (DMSO, TMP-Cl) incubated in ENR and NR media, respectively, for **(D-H)**; n = 3 for Western Blot in **J**. Scale bar = 50 μm.

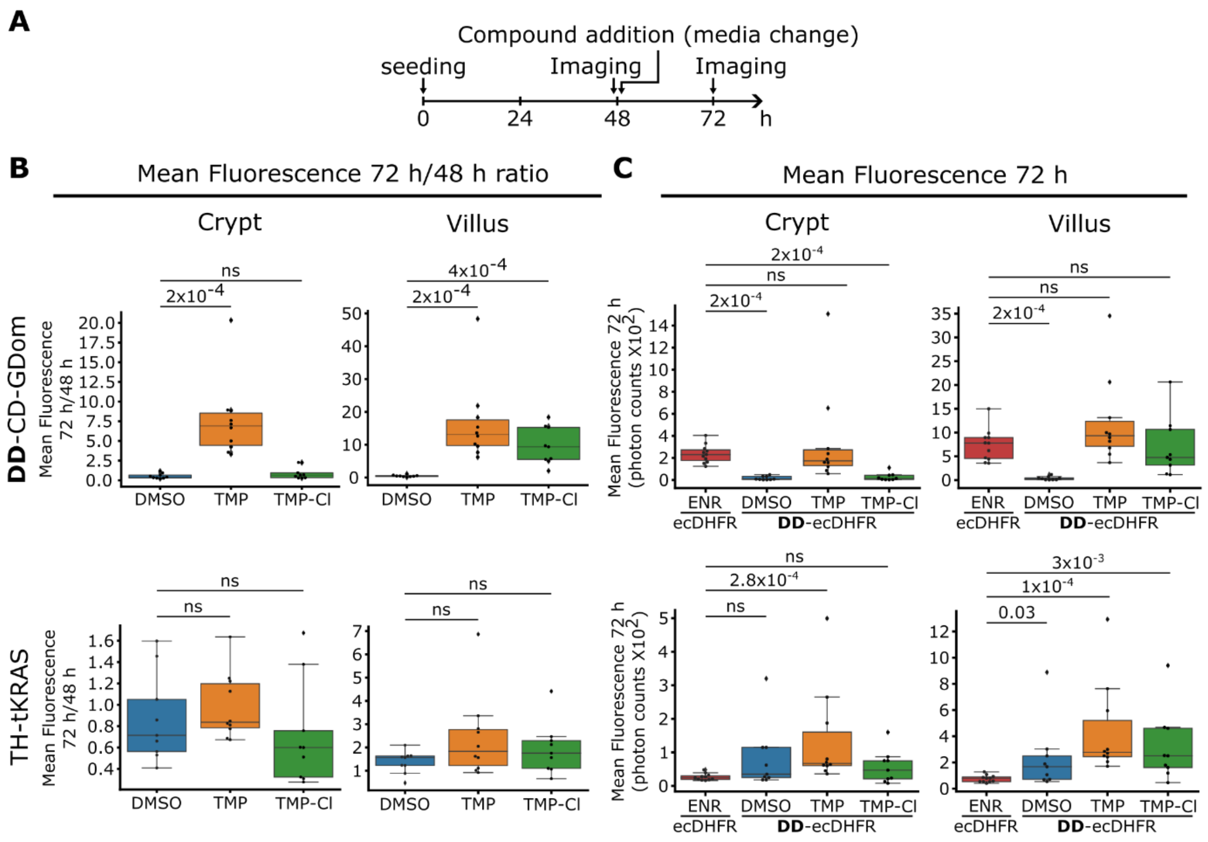


**Figure S6. Stabilization of DD-CD-GDom in crypt and villus subregions in Opto-CID-DD-KRASG12V-mSIOs.**

**(A)** Scheme depicting the time for seeding, imaging, and compound addition for all experiments. **(B)** Boxplots of the fluorescence mean intensity 24 h after / 1 h before compound addition ratio from crypts and villus regions (left and right, respectively), labeled as the time after mSIOs individual crypts were seeded (72 h / 48 h). **(C)** Boxplot comparing the mean fluorescence intensity of Opto-CID-DD-KRASG12V-mSIOs 24 h after compound addition (72 h after seeding) with Opto-CID-KRASG12V-mSIOs bearing the stable ecDHFR domain. n = 9, 10, and 9 for DMSO, TMP, and TMP-Cl Opto-CID-DD-KRASG12V-mSIOs and 11 for Opto-CID-KRASG12V-mSIOs. *p*-value was calculated by the Mann-Whitney U test.
